# RANKL from Bone Marrow Adipose Lineage Cells Promotes Osteoclast Formation and Bone Loss

**DOI:** 10.1101/2020.09.12.294348

**Authors:** Yan Hu, Xiaoqun Li, Xin Zhi, Wei Cong, Biaotong Huang, Huiwen Chen, Yajun Wang, Yinghua Li, Lipeng Wang, Chao Fang, Jiawei Guo, Ying Liu, Jin Cui, Liehu Cao, Weizong Weng, Qirong Zhou, Sicheng Wang, Xiao Chen, Jiacan Su

## Abstract

Receptor activator of NF-κB ligand (RANKL) is essential for osteoclast formation and bone remodeling. Nevertheless, the cellular source of RANKL for osteoclastogenesis has not been fully uncovered. Different from peripheral adipose tissue, bone marrow (BM) adipose lineage cells originate from bone marrow mesenchymal stromal cells(BMSCs). Here we demonstrate that adiponectin promoter-driven Cre expression (*Adipoq^Cre^*) can target bone marrow adipose lineage cells. We cross the *Adipoq^Cre^* mice with *rankl^fl/fl^* mice to conditionally delete RANKL from BM adipose lineage cells. Conditional deletion of RANKL increases cancellous bone mass of long bones in mice by reducing the formation of trabecular osteoclasts and inhibiting bone resorption but does not affect cortical bone thickness or resorption of calcified cartilage. *Adipoq^Cre^; rankl^fl/fl^* mice exhibit resistance to estrogen deficiency and rosiglitazone (ROS) induced trabecular bone loss but show bone loss induced by unloading. BM adipose lineage cells therefore represent an essential source of RANKL for the formation of trabecula osteoclasts and resorption of cancellous bone during remodeling under physiological and pathological conditions. Targeting bone marrow adiposity is a promising way of preventing pathological bone loss.

## Introduction

Osteoclasts are specialized multinucleated cells derived from the monocyte-macrophage hematopoietic lineage and are responsible for resorption of bone matrix (Boyle *et al*, 2003). During longitudinal bone growth, hypertrophic chondrocytes attract osteoclasts and blood vessels to direct the matrix mineralization (Kronenberg, 2003). Under normal conditions, trabecular bone is resorbed periodically by osteoclasts followed by osteogenesis in the cavities, in a process known as remodeling (Raggatt & Partridge, 2010). Excess bone resorption by osteoclasts results in pathological bone loss disorders, including postmenopausal osteoporosis, Paget’s disease, and others (Manolagas & Jilka, 1995).

Receptor activator of NF-κB ligand (RANKL) is an essential mediator of osteoclast formation during bone resorption (Sobacchi *et al*, 2007). Matrix-embedded hypertrophic chondrocytes and osteocytes serve as the primary sources of RANKL for mineralized cartilage resorption in the development of growing bone and remodeling of adult bone, respectively (Xiong *et al*, 2011). Initially RANKL is produced in a membranous form that can be cleaved by proteases to produce soluble RANKL (Lacey *et al*, 1998). Because most matrix-embedded cells do not come in direct contact with osteoclast progenitors, these cells may control osteoclast formation through the production of soluble RANKL (Liu *et al*, 2001). Although it does contribute to osteoclast formation and the resorption of cancellous bone in adult bone, soluble RANKL does not affect cancellous bone mass or structure in growing bone (Xiong *et al*, 2018). Moreover, bone loss caused by estrogen deficiency in ovariectomized mice is not prevented or reduced by a lack of soluble RANKL (Xiong *et al*., 2018). These results indicate that the membrane-bound form of RANKL is sufficient for regulating osteoclast formation in a direct contact manner. As RANKL is expressed in various cell types, the cellular source of RANKL for osteoclastogenesis during bone growth and estrogen deficiency remains unclear.

Unlike white or brown adipose tissue, bone marrow (BM) adipose lineage cells derived from BM mesenchymal stromal cells (BMSCs) by lineage tracing have been implicated in osteoclastogenesis and hematopoiesis (Zhou *et al*, 2017; Zhou *et al*, 2014). The number of BM adipose lineage cells increases with age as well as under certain pathological conditions (Duque *et al*, 2011). BM adipose lineage cells express RANKL, a phenotype not seen in either white or brown adipose tissue (Fan *et al*, 2017), and increased BM adipose tissue is associated with increased osteoclast formation (Yu *et al*, 2018). Bone marrow-derived preadipose cell line MC3T3-G2/PA6 cells support osteoclast formation in the co-culture system in the presence of 1α,25-(OH)_2_D3 and dexamethasone (N. *et al*, 1989). Based on these observations, we hypothesized that BM adipose lineage cells could serve as a source of RANKL for osteoclast formation and pathological bone resorption. We conditionally deleted RANKL in BM adipose lineage cells by crossing the *Adipoq^Cre^* mice with *rankl^fl/fl^* mice. We found that RANKL derived from BM adipose lineage cells did not affect resorption of calcified cartilage in the growth plate or cortical bone but did contribute to osteoclast formation and resorption of the cancellous bone in the longitudinal bone. Mice lacking RANKL in BM adipose lineage cells were protected from bone loss after ovariectomy or rosiglitazone administration but not from bone loss induced by unloading. BM adipose lineage cells therefore represent an essential source of RANKL for the trabecular osteoclast formation and cancellous bone resorption in physiological and pathological bone resorption.

## Results

### 1. Conditional knockout of RANKL in bone marrow adipose lineage cells

As previously reported (Ambrosi *et al*, 2017; Wang *et al*, 2020b), we used the *Adipoq^Cre^* to specifically delete RANKL in BM adipose lineage cells. BMSCs consist of three sub-populations: multi-potent stem cell-like population, osteochondrogenic progenitor cell (OPC) and adipogenic progenitor cell (APC), while APC will irreversibly mature toward a preAd stage: pre-adipocyte (preAd) (Ambrosi *et al*., 2017). To identify the subgroup targeted by the *Adipoq^Cre^*, we crossed *Adipoq^Cre^* mice with *Rosa26-lsl-tdTomato* mice to generate *Adipoq^Cre^; R26^tdTomato^* mice, which permits identification of Adipoq-Cre targeted cells. Flow cytometry analysis of the femoral bone marrow showed that CD45^−^CD31^−^Sca-1^+^ cells representing APC (CD45^−^CD31^−^ Sca-1^+^CD24^−^) and multi-potent stem cells (CD45^−^CD31^−^Sca-1^+^CD24^+^) did not express tdTomato (Fig 1A). Among CD45^−^CD31^−^Sca-1^−^ cells covering preAd (CD45^−^ CD31^−^Sca1^−^Reep2^+^) and OPC (CD45^−^CD31^−^Sca1^−^PDGFα^+^), tdTomato^+^Reep2^+^ preAd accounted for 16.0% and tdTomato^+^PDGFα^+^ OPC accounted for 0.186%. The flowcytometry results indicated that the *Adipoq^Cre^* targeted preAd in bone marrow, which is consistent with a recent study by Qin, et al.(Zhong *et al*, 2020). The immunofluorescence staining of Aggrecan and DAPI showed that the *Adipoq^Cre^* did not elicit any reporter gene activity in osteocytes (Fig 1Bi) or chondrocytes (Fig 1Bii). To further verify the specificity of the Cre line, we sorted tdTomato^+^ cells from *Adipoq^Cre^; R26^tdTomato^* mice and conducted proliferation and differentiation assays *in vitro* and *in vivo*. The Giemsa staining showed that the BMSCs from control wildtype mice could form CFU-F colonies *in vitro*, while the tdTomato^+^ cells attached to the culture dish but could not form CFU-F colonies (Fig EV1A-B). Then the adipogenic and osteogenic potential of BMSCs and tdTomato^+^ cells were assessed by oil red O staining and Alizarin red staining, respectively. The results showed that BMSCs could differentiate into osteoblasts and adipocytes *in vitro*, while tdTomato^+^ cells could only undergo adipogenesis (Fig EV1C, E). The qPCR results of peroxisome proliferator-activated receptor γ (PPARγ), lipoprotein lipase (LPL), osteocalcin (OCN) and alkaline phosphatase (ALP) showed consistent results with differentiation assays (Fig EV1D, F). For *in vivo* assay, we injected BMSCs and tdTomato^+^ cells into mice kidney capsules which were harvested after 4 weeks. Immunofluorescence staining of OCN, Aggrecan and Perilipin was performed. Results indicated that BMSCs formed osteoblasts, adipocytes and chondrocytes after transplanted and sorted tdTomato^+^ cells from *Adipoq^Cre^; R26^tdTomato^* mice only formed adipocytes (Fig EV2A-C).

**Figure 1.**
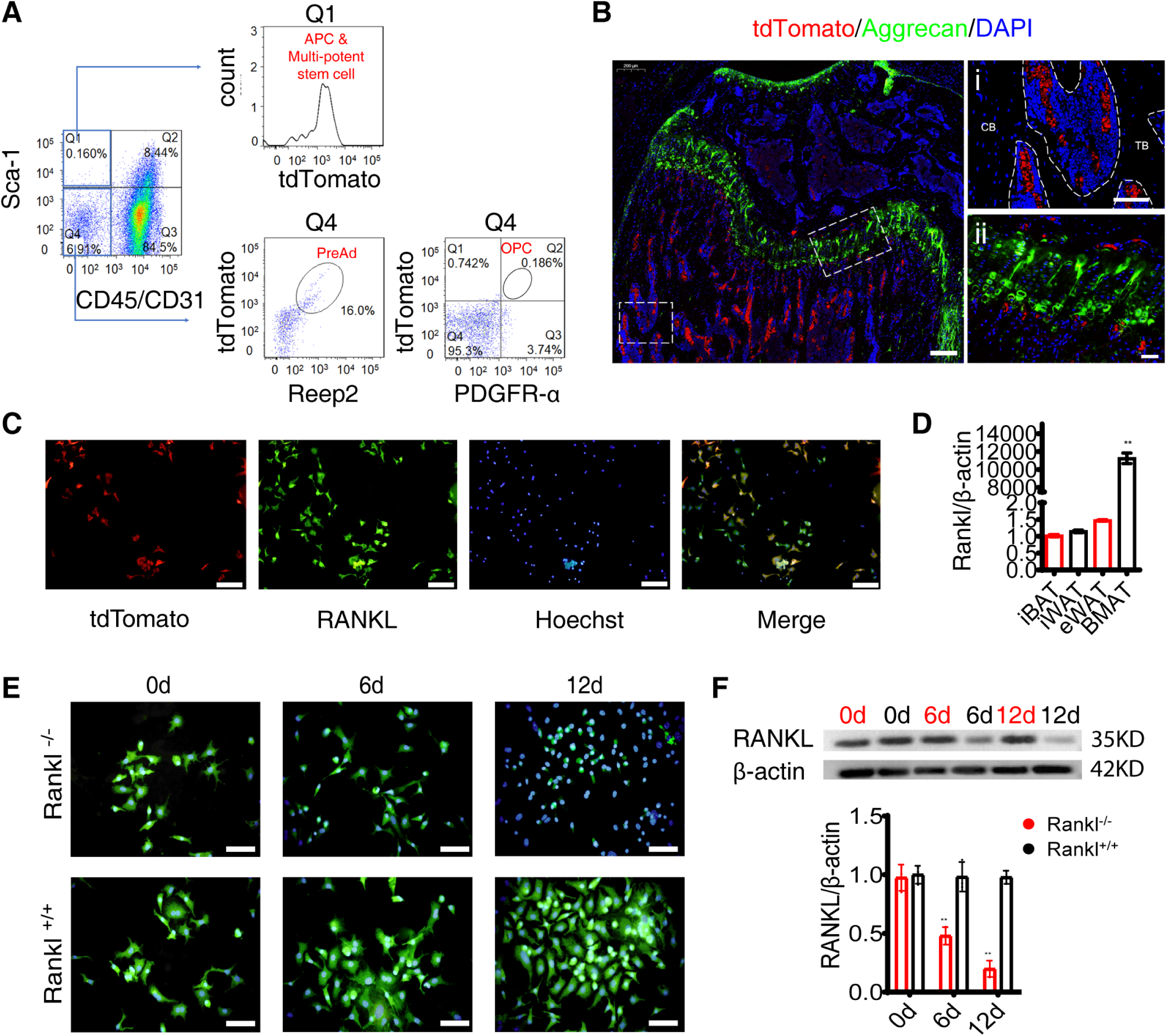
*Adipoq^Cre^* targets BM adipose lineage cells. (A) FACS analysis of tdTomato^+^ cells in the bone marrow cells (percentages represent average values). (B) Immunofluorescence staining of aggrecan and DAPI of femur sections from *Adipoq^Cre^; R26^tdTomato^* mice. Scale bar=200/100/50µm in left/ upper right/ bottom right images. CB: cortical bone, TB: trabecular bone. (C) Immunofluorescence staining of RANKL and Hoechst in bone marrow cells from *Adipoq^Cre^; R26^tdTomato^* mice. Scale bar=50 µm. (D) qPCR results of *Rankl* in BMAT and peripheral adipose tissues, including epididymal (eWAT), inguinal (iWAT), and interscapular (iBAT) (n=3 independent biological replicates). Data were compared using one-way ANOVA (** indicates P < 0.01), error bars are standard deviations. (E) Immunofluorescence staining of RANKL during *in vitro* adipogenesis of BMSCs from *Adipoq^Cre^; rankl^fl/fl^* mice (*Rankl^-/-^*) and *rankl^fl/fl^* mice (*Rankl^+/+^*). Scale bar=50 μm. (F) Western blotting analyses of RANKL in BMSCs during *in vitro* BMSCs adipogenesis (n=3 independent biological replicates). Data were compared using an unpaired t-test (** indicates P < 0.01), error bars are standard deviations.

In isolated bone marrow cells from *Adipoq^Cre^; R26^tdTomato^* mice, immunofluorescent staining showed that RANKL colocalized with all tdTomato^+^ cells, which indicated that all BM adipose lineage cells expressed RANKL (Fig 1C). To verify the specificity of RANKL expression in BM adipose tissue, we examined the *rankl* gene transcription in epididymal adipose tissue (eWAT), inguinal adipose tissue (iWAT), interscapular brown adipose tissue (iBAT) and BM adipose tissue (BMAT) from wild type C57BJ/6L mice by qPCR as previous reported.(Fan *et al*., 2017) The results showed that different from peripheral adipocytes, the BM adipose lineage cells highly expressed *rankl* (Fig 1D). Then we isolated BMSCs from *Adipoq^Cre^; rankl^fl/fl^* mice and induced adipogenic differentiation. During adipogenesis, expression of RANKL in BMSCs decreased markedly in *Adipoq^Cre^; rankl^fl/fl^* mice but remained unchanged in *rankl^fl/fl^* mice (Fig 1E-F). These results demonstrate that the *Adipoq^Cre^* efficiently deletes RANKL in bone marrow adipose lineage cells and but not in chondrocytes, osteocytes or mesenchymal progenitors.

### 2. RANKL knockout from BM adipose lineage cells increases trabecular bone mass

To assess the role of BM adipocyte RANKL in bone mass, we established BM adipocyte RANKL knockout mice by crossing the *Adipoq^Cre^* mice with *rankl^fl/fl^* mice (Appendix Fig S1A). Both male and female mice were used in the study. Conclusions are based on data analyses from both sexes, although only data from females are presented here. For both male and female mice, the body weight and body length were recorded from birth to 8 weeks (Appendix Fig S1B). And *Adipoq^Cre^; rankl^fl/fl^* mice showed a slightly increase of body weight compared to their control littermates. RANKL deletion in BM adipose lineage cells did not affect RANKL expression in spleen (Appendix Fig S1C) by immunohistochemistry, spleen (Appendix Fig S1D) and lymph node (Appendix Fig S1E) development by HE staining, or differentiation of T and B cells by flowcytometry (Appendix Fig S1F-G).

We harvested the femurs of *Adipoq^Cre^; rankl^fl/fl^* and *rankl^fl/fl^* mice at 4, 8 and 16 weeks. Quantitative computed tomography (μ-QCT) analyses revealed significant increases in trabecular bone mass of *Adipoq^Cre^; rankl^fl/fl^* mice compared to *rankl^fl/fl^* mice as confirmed by increased bone mineral density (BMD), trabecular bone volume (BV/TV), trabecular number (Tb.N) and decreased trabecular spacing (Tb.Sp) (Fig 2A-B). No differences in cortical bone thickness or BMD were found between the two groups at 8 weeks (Fig 2C-D). Taken together, these results indicate that deleting RANKL from BM adipose lineage cells increase trabecular bone mass.

**Figure 2.**
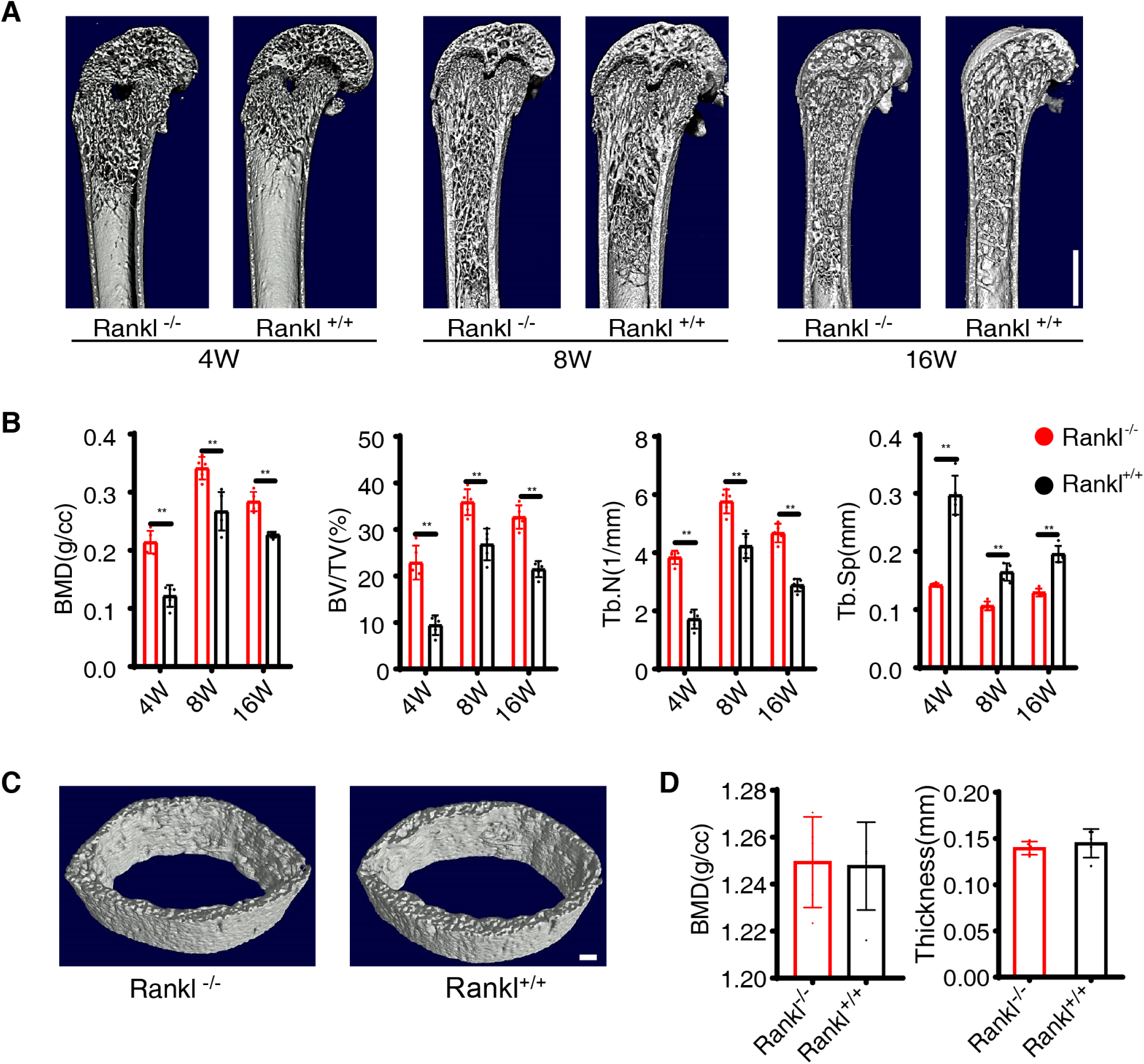
Cancellous bone mass increases in *Adipoq^Cre^; rankl^fl/fl^* mice. (A-B) Micro-CT and statistical analyses of BMD, BV/TV, Tb.N and Tb.Sp in trabecular bone from *Adipoq^Cre^; rankl^fl/fl^* (*Rankl^-/-^*) and *rankl^fl/fl^* (*Rankl^+/+^*) mice at 4 weeks, 8 weeks and 16 weeks (n=5 independent biological replicates). Scale bar=1 mm. Data were compared using an unpaired t-test (** indicates P < 0.01), error bars are standard deviations. (C) Micro-CT images of cortical bone from *Rankl^-/-^* and *Rankl^+/+^* mice at 8 weeks. Images are representative of five independent biological replicates. Scale bar =1 mm. (D) Micro-CT analyses and statistical analyses of cortical bone from *Rankl^-/-^* and *Rankl^+/+^* mice at 8 weeks (n=5 independent biological replicates). Data were compared using an unpaired t-test, error bars are standard deviations.

### 3. RANKL from BM adipose lineage cells is essential for trabecular bone remodeling

Tartrate-resistant acid phosphatase (TRAP) staining and OCN immunofluorescence staining was performed to assess the roles of BM adipocyte-derived RANKL in trabecular bone remodeling. TRAP staining of trabecular bone revealed a significant decrease in the number of TRAP^+^ cells (osteoclasts) in the bone marrow around the trabecular bones of *Adipoq^Cre^; rankl^fl/fl^* mice compared to *rankl^fl/fl^* mice at 8 weeks (Fig 3A a1 vs. a4, B). The number of TRAP^+^ cells under the growth plate (Fig 3A a2 vs. a5, C) and periosteal TRAP^+^ cells (Fig 3A a3 vs. a6, D) did not show significant difference. The thickness of growth plate cartilage (Fig EV3A) and tooth eruption (Fig EV3B) were not significantly different between two groups.

**Figure 3.**
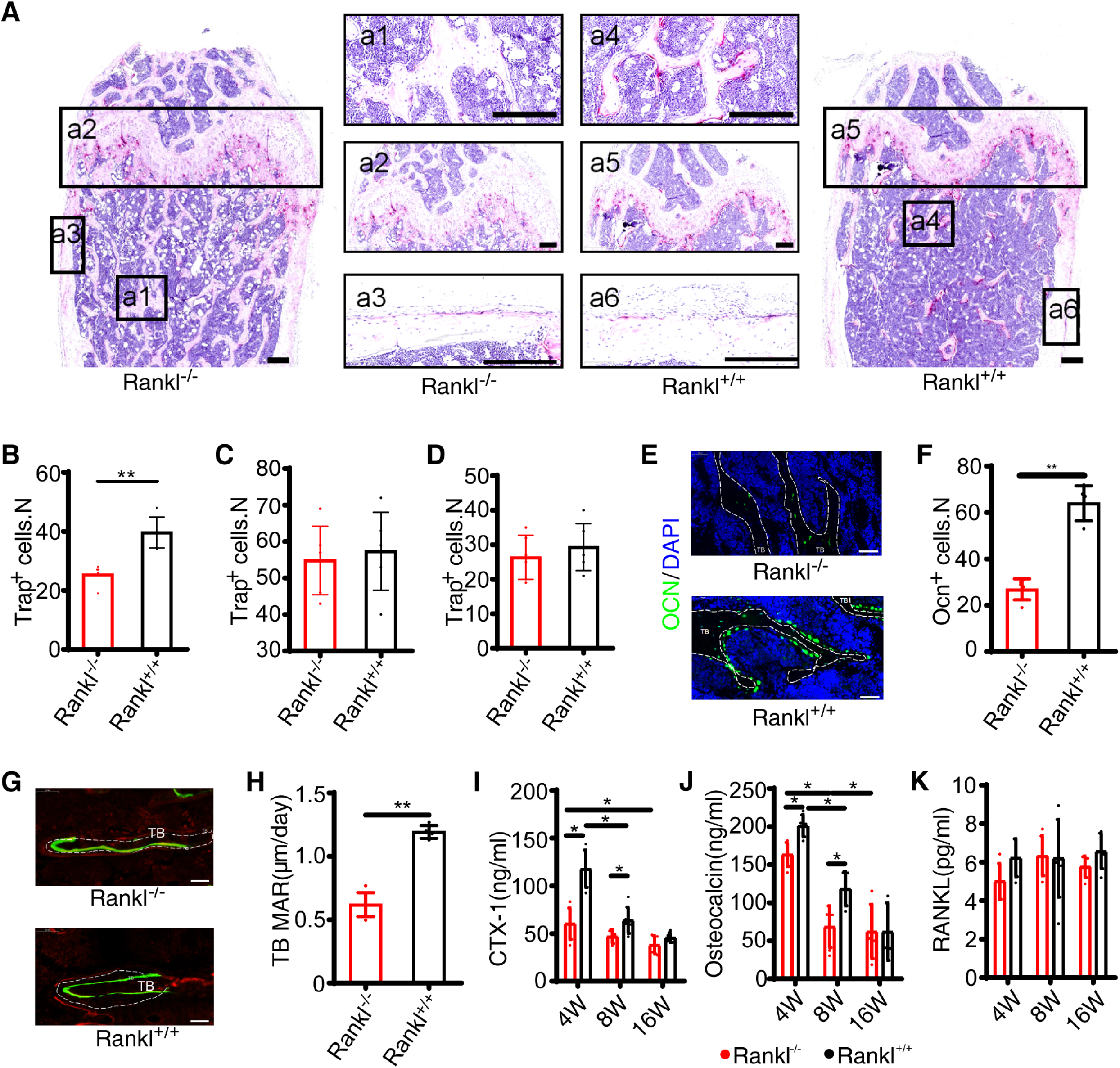
RANKL knockout from BM adipose lineage cells regulates trabecular bone remodeling. (A) Representative images of TRAP staining of femur from *Adipoq^Cre^; rankl^fl/fl^* (*Rankl^-/-^*) and *rankl^fl/fl^* (*Rankl^+/+^*) mice at 8 weeks. Scale bar=200 μm. Panel a1 and a4 indicate trabecular bone, a2 and a5 stand for growth plate, a3 and a6 denote cortical bone. (B) Numbers of Trap^+^ cells on trabecular bone (a1 vs. a4, see (A)) (n=6 independent microscopic vision fields from random mice samples). Data were compared using an unpaired t-test (** indicates *P* < 0.01), error bars are standard deviations. (C) Numbers of Trap^+^ cells under the growth plate (a2 vs. a5, see (A)) (n=6 independent microscopic vision fields from random mice samples). Data were compared using an unpaired t-test, error bars are standard deviations. (D) Numbers of periosteal Trap^+^ cells (a3 vs. a6, see (A)) (n=6 independent microscopic vision fields from random mice samples). Data were compared using an unpaired t-test, error bars are standard deviations. (E) Immunofluorescence staining of osteocalcin from 8-week-old mice (n=6 mice in *Rankl^-/-^* group and 4 mice in *Rankl^+/+^* group). Scale bar=50 μm. TB: trabecular bone. (F) Statistical analyses of Ocn^+^ cells (n=6 independent microscopic vision fields from random mice samples). Data were compared using an unpaired t - test (** indicates P < 0.01), error bars are standard deviations. (G) Calcein staining of trabecular bone at 8 weeks. Scale bar=50 μm. TB: trabecular bone. (** indicates *P* < 0.01), error bars are standard deviations. (H) Mineral apposition rate (MAR) of trabecular bone at 8 weeks.(n=6 mice in *Rankl*^-/-^ group and 5 mice in *Rankl*^+/+^ group). Data were compared using an unpaired t-test (** indicates P < 0.01), error bars are standard deviations. (I-K) Serum CTX-1, OCN and bone marrow RANKL levels from *Rankl^-/-^* and *Rankl^+/+^* mice at 4 weeks (n=5 and 5), 8 weeks (n=5 and 6) and 16 weeks (n=5 and 5). Data were compared using an unpaired t-test (* indicates *P* < 0.05, ** indicates *P* < 0.01), error bars are standard deviations.

OCN immunofluorescence staining showed fewer osteoblasts around the trabeculae in *Adipoq^Cre^; rankl^fl/fl^* mice than in *rankl^fl/fl^* mice at 8 weeks (Fig 3E-F). Calcein staining revealed less trabecular bone formation and lower trabecular mineralization apposition rate (MAR) in *Adipoq^Cre^; rankl^fl/fl^* mice relative to *rankl^fl/fl^* mice at 8 weeks (Fig 3G-H). By contrast, the endosteal and periosteal bone formation was not significantly different between two groups (Fig EV3C-D). The bone histomorphometric parameters of *Adipoq^Cre^; rankl^fl/fl^* mice and *rankl^fl/fl^* mice were showed in Appendix Table S1. The above results suggested that RANKL deletion from BM adipose lineage cells impeded trabecular osteoclasts formation and the following osteogenesis and bone formation, namely bone remodeling.

We isolated BMSCs and *in vitro* Alizarin red staining after osteogenic differentiation showed no significant difference between the two groups (Appendix Fig S2A). Expression of alkaline phosphatase (ALP) and OCN was comparable in *Adipoq^Cre^; rankl^fl/fl^* and *rankl^fl/fl^* mice (Appendix Fig S2B-C). *In vitro* oil red O staining and expression of LPL and PPARγ after adipogenic differentiation of BMSCs revealed no significant difference between *Adipoq^Cre^; rankl^fl/fl^* and *rankl^fl/fl^* mice (Appendix Fig S2D-F). The above results indicated that RANKL deletion from BM adipose lineage cells did not affect BM osteogenesis and adipogenesis.

Next, we examined serum CTX-1, OCN levels and bone marrow RANKL levels. Compared to *rankl^fl/fl^* mice, *Adipoq^Cre^; rankl^fl/fl^* mice had lower serum CTX-1 at 4, 8 and 16 weeks (Fig 3I) and lower serum OCN at 4 and 8 weeks (Fig 3J). Overall, both serum CTX-1 and OCN decreased with age (Fig 3I-J). Bone marrow RANKL levels were not significantly different between *Adipoq^Cre^; rankl^fl/fl^* and *rankl^fl/fl^* mice at 4, 8 and 16 weeks (Fig 3K), indicating that soluble RANKL level was not affected by RANKL knockout in BM adipose lineage cells. The above results indicated that BM adipocyte RANKL knockout decreased trabecular bone remodeling but did not affect the periosteal bone formation, calcified cartilage resorption at the growth plate.

A recent study reported that the *Adipoq^Cre^* targets PDGFR^+^VCAM-1^+^ stromal cells,(Mukohira *et al*, 2019) and our previous work showed that RANKL signaling in BMSCs negatively regulates osteoblastogenesis and bone formation (Chen *et al*, 2018a). To rule out the effects of RANKL signaling on osteoblast differentiation of BMSCs *in vivo*, we generated the *Adipoq^Cre^; rank^fl/fl^* conditional knockout mice. Micro-CT analyses revealed no significant differences between *rank* knockout in BM adipose lineage cells and *rank^fl/fl^* mice at any time point tested (Fig EV4A-B). Overall, these results indicate that BM adipose lineage cells are an essential source of RANKL for trabecular bone osteoclast formation and subsequent remodeling.

### 4. RANKL from BM adipose lineage cells does not mediate unloading-induced bone resorption

Bone constantly adapts its structure in response to mechanical signals under the control of osteoblastic cells, which senses mechanical loading and regulates osteoclasts formation and trabecular bone remodeling through RANKL (Xiong *et al*., 2011). To further determine if BM adipocyte derived RANKL mediates skeletal mechanical loading, we carried out unloading test by tail suspension in 8-week-old mice for 4 weeks. In each group, micro-CT results showed the trabecular bone BMD, BV/TV and Tb.N were significantly decreased and the Tb.Sp was increased (Fig 4A-B). The cortical changes were consistent with trabecular bones (Fig 4C-D). The *Adipoq^Cre^; rankl^fl/fl^* mice did not prevent bone loss induced by tail suspension. The results support that BM adipocyte RANKL does not mediate unloading-induced bone resorption.

**Figure 4.**
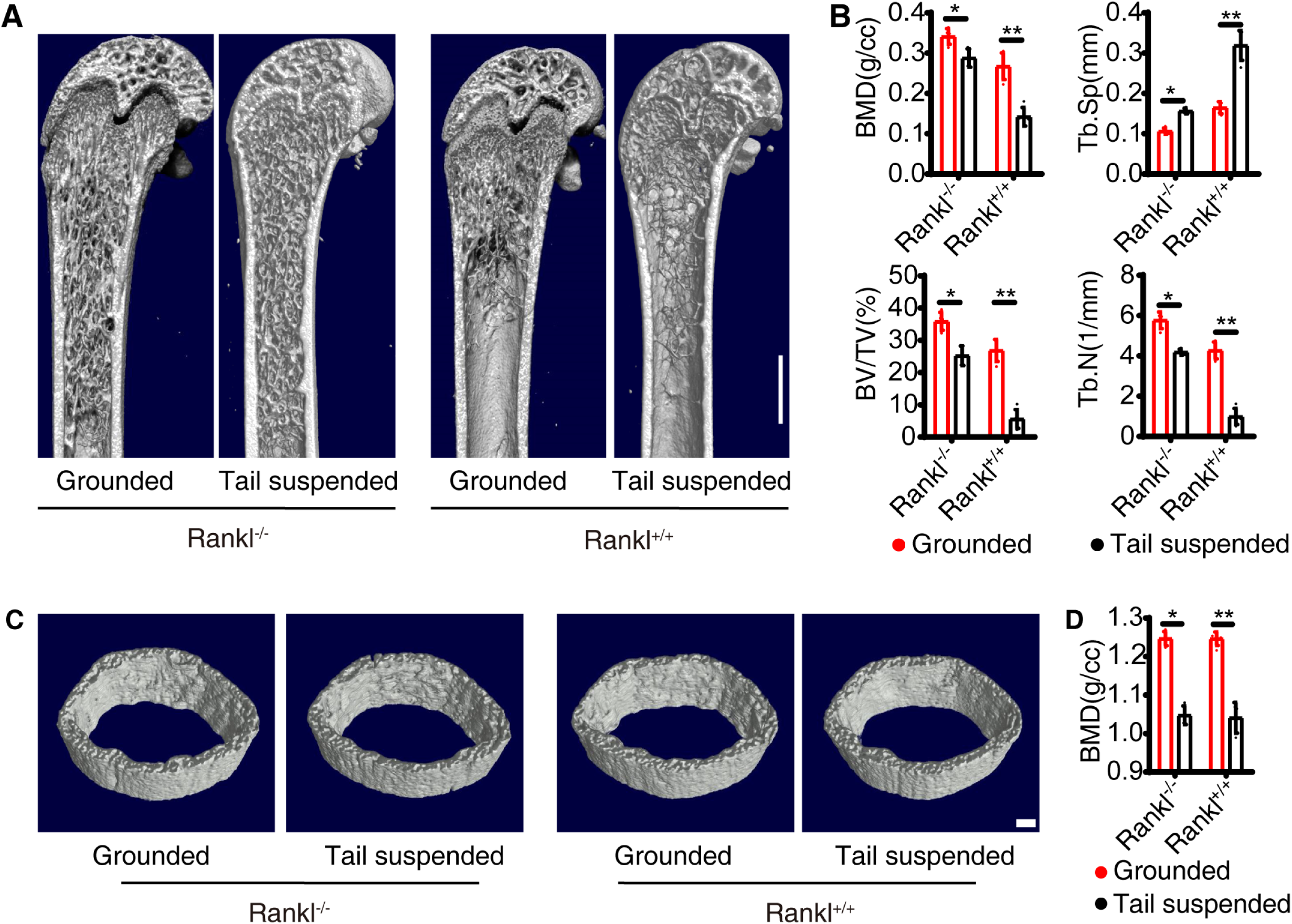
RANKL knockout from BM adipose lineage cells does not affect osteocyte regulation on osteoclasts. (A-B) Micro-CT analyses of trabecular bone from Grounded and Tail suspended model (n=5 independent biological replicates). Scale bar=1 mm. Data were compared using an unpaired t-test (* indicates *P* < 0.05, ** indicates *P* < 0.01), error bars are standard deviations. (C-D) Micro-CT analyses of cortical bone (n = 5 independent biological replicates). Scale bar=1 mm. Data were compared using an unpaired t-test (* indicates P < 0.05, ** indicates *P* < 0.01), error bars are standard deviations..

### 5. RANKL from BM adipose lineage cells mediates osteoclastogenesis and bone loss in ovariectomized mice

Increased BM adipose tissue is associated with increased fracture risk in postmenopausal osteoporosis (Veldhuis-Vlug & Rosen, 2018). To examine the roles of RANKL from BM adipose lineage cells in bone resorption after estrogen withdrawal, we used a mouse model of ovariectomy (OVX)-induced bone loss. Immunofluorescence staining of adiponectin revealed a significant increase in the number of BM adipose lineage cells following OVX in both *Adipoq^Cre^; rankl^fl/fl^* and *rankl^fl/fl^* mice, with no statistical difference between groups (Fig EV5A-B). At 6 weeks post-ovariectomy, trabecular bone mass was significantly decreased in *rankl^fl/fl^* mice, a phenotype not seen in *Adipoq^Cre^; rankl^fl/fl^* mice (Fig 5A-B). Consistent results were evident via H&E staining (Fig EV5C). TRAP staining showed that in *rankl^fl/fl^* mice, after OVX the number of osteoclasts significantly increased while in *Adipoq^Cre^; rankl^fl/fl^* mice the number did not significantly change (Fig 5C-D). The OCN staining for osteoblasts demonstrated consistent results with TRAP staining (Fig 5E-F). These results indicated that increased bone remodeling after OVX was inhibited in *Adipoq^Cre^; rankl^fl/fl^* mice compared to their littermates. ELISA analyses revealed increased serum CTX-1 (Fig EV5D), OCN (Fig EV5E) and bone marrow RANKL (Fig EV5F) levels in *rankl^fl/fl^* OVX mice relative to non-OVX controls, with no significant changes in protein abundance in *Adipoq^Cre^; rankl^fl/fl^* OVX mice. These results demonstrate that RANKL from BM adipose lineage cells is an essential mediator of excess osteoclasts formation, bone resorption and increased bone turnover after estrogen withdrawal.

**Figure 5.**
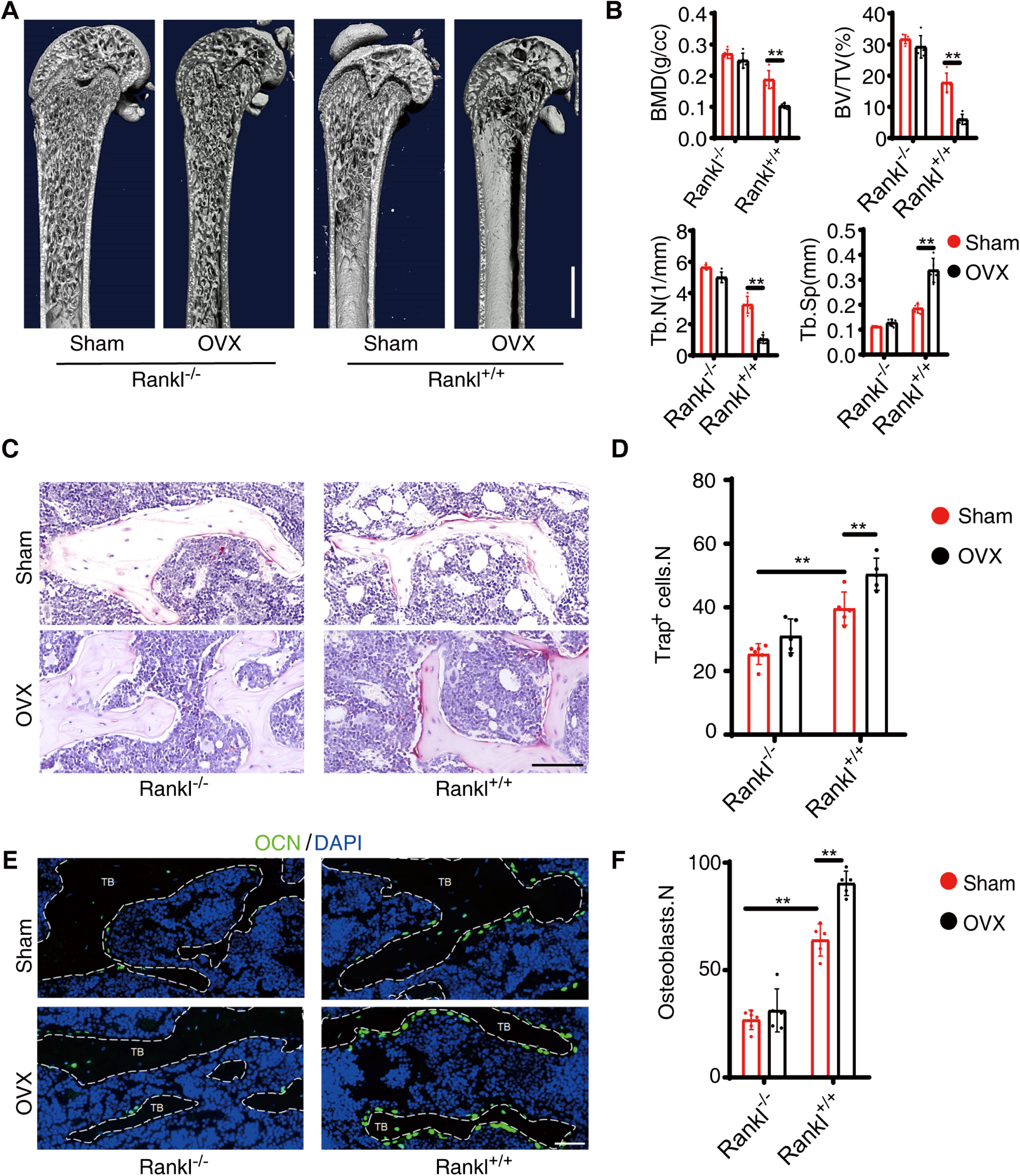
RANKL knockout from BM adipose lineage cells contributes to excess osteoclast formation and bone resorption in OVX mice. (A-B) Micro-CT analyses of sham and OVX model of *Adipoq^Cre^; rankl^fl/fl^* (*Rankl^-/-^*) and *rankl^fl/fl^* (*Rankl^+/+^*) mice (n=5 independent biological replicates). Scale bar=1 mm. Data were compared using an unpaired t-test (** indicates *P* < 0.01), error bars are standard deviations. (C-D) TRAP staining and statistical analyses of numbers of osteoclasts in Sham and OVX model (Images are representative of 5 independent biological replicates in (C), and n=5 independent microscopic vision fields in (D)). Scale bar=100 µm. Data were compared using an unpaired t-test (** indicates *P* < 0.01), error bars are standard deviations. (E-F) Immunofluorescence staining of OCN and statistical analyses of the number of osteoblasts (Images are representative of 5 independent biological replicates in (E), and n=5 independent microscopic vision fields in (F)). Scale bar=50 µm. TB: trabecular bone. Data were compared using an unpaired t - test (** indicates *P* < 0.01), error bars are standard deviations.

### 6. RANKL from BM adipose lineage cells contributes to rosiglitazone-induced osteoclastogenesis and bone loss

Rosiglitazone (ROS), a PPARγ activator used to treat type 2 diabetes, is associated with increased BM adipogenesis and fracture risk in both men and women (Aubert *et al*, 2010; Kahn *et al*, 2008). The mechanisms underlying these effects are still poorly understood but are likely attributable to increased adipogenesis due to PPARγ activation (Wei *et al*, 2010). To explore the roles of BM adipocyte RANKL in this process, we used a model of ROS-induced bone loss. In this model, ROS (10 mg/kg) was administered for 6 weeks. Adiponectin immunofluorescent staining revealed a substantial increase in the number of BM adipose lineage cells after ROS administration in both *Adipoq^Cre^; rankl^fl/fl^* and *rankl^fl/fl^* mice, with no significant differences between the groups (Appendix Fig S3A-B). Trabecular bone mass showed a significant decrease in *rankl^fl/fl^* mice but no decrease in *Adipoq^Cre^; rankl^fl/fl^* mice (Fig 6A-B). H&E staining showed consistent results with CT analyses (Appendix Fig S3C). TRAP staining showed that ROS treatment significantly increased the number of osteoclasts in *rankl^fl/fl^* mice but not in *Adipoq^Cre^; rankl^fl/fl^* mice (Fig 6C-D). OCN immunofluorescent staining showed that the number of osteoblasts was significantly increased in *rankl^fl/fl^* mice after ROS treatment but not in *Adipoq^Cre^; rankl^fl/fl^* mice (Fig 6E-F). Serum CTX-1 (Appendix Fig S3D) and OCN levels were increased in response to ROS in *rankl^fl/fl^* mice (Appendix Fig S3E) but not in *Adipoq^Cre^; rankl^fl/fl^* mice. BM RANKL levels showed no significant differences in *Adipoq^Cre^; rankl^fl/fl^* mice after treated with ROS (Appendix Fig S3F). These results indicate that increased BM adipose lineage cells are an important source of RANKL for osteoclast formation and bone loss after ROS administration.

**Figure 6.**
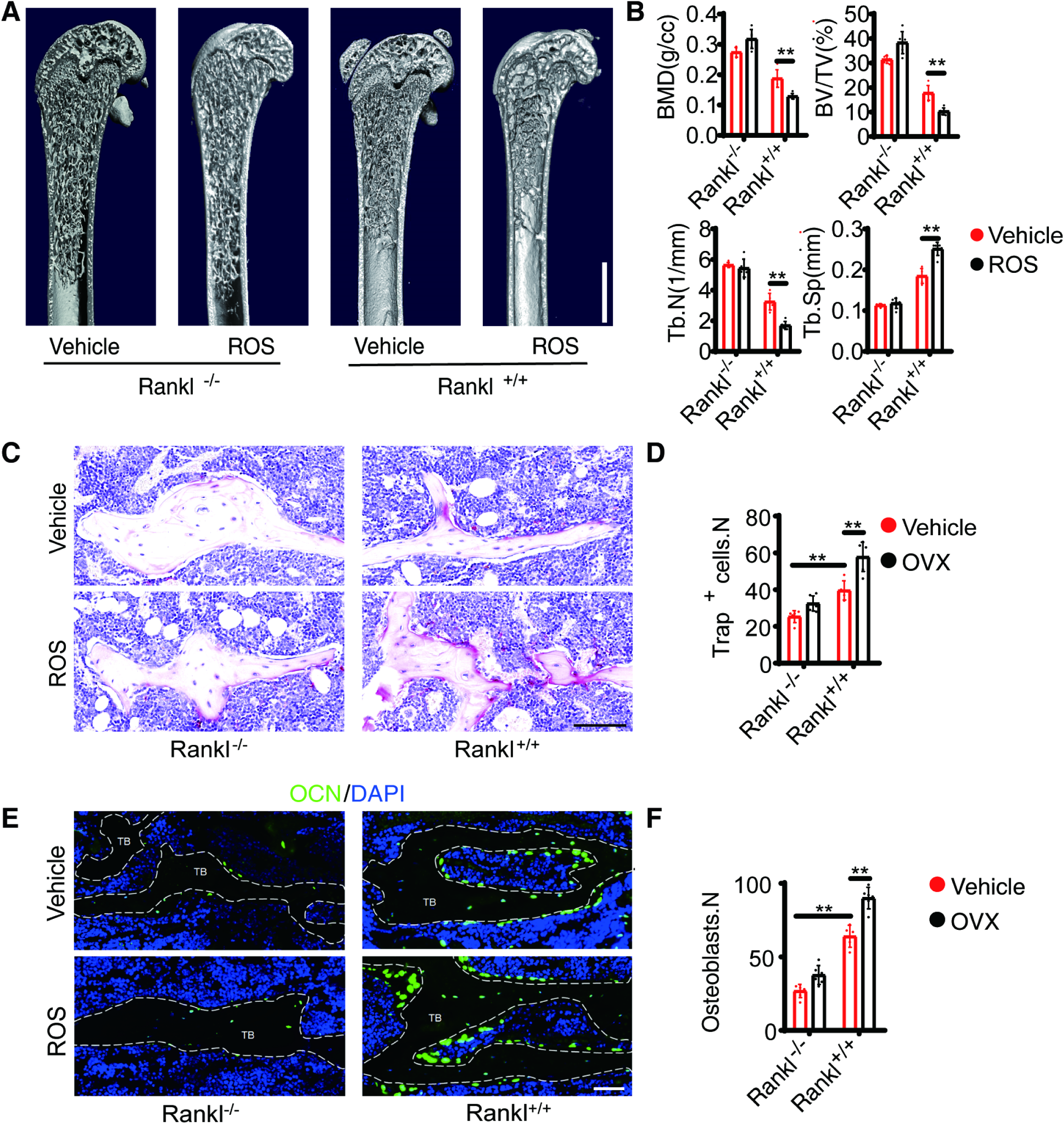
RANKL knockout from BM adipose lineage cells mediates rosiglitazone-induced bone loss. (A-B) Three-dimensional CT analyses of BMD, BV/TV, Tb.N, and Tb.Sp in trabecular bone from Vehicle (n=5 independent biological replicates) and ROS (n=6 independent biological replicates) model of *Adipoq^Cre^; rankl^fl/fl^* (*Rankl^-/-^*) and *rankl^fl/fl^* (*Rankl^+/+^*) mice. Scale bar=1 mm. indicates *P* < 0.01), error bars are standard deviations. (C-D) TRAP staining and statistical analyses of numbers of osteoclasts in the trabecular bone from Vehicle (n=6 and 5 independent biological replicates) and ROS (n=6 and 6 independent biological replicates) treatment. Scale bar=100 µm. Data standard deviations. n=5 independent microscopic vision fields in (D). (E-F) OCN staining and statistical analyses of the number of osteoblasts in femurs from Vehicle (n=6 and 5 independent biological replicates) and ROS (n=6 and 6 independent biological replicates) treatment. Scale bar=50 µm. TB: trabecular bone. Data were compared using an unpaired t-test (** indicates *P* < 0.01), error bars are standard deviations. n=5 independent microscopic vision fields in (F).

## Discussion

As an essential factor for osteoclastogenesis, RANKL is widely expressed by a variety of cells types within the bone marrow (Nakashima *et al*, 2011). Factors including parathyroid hormone, mechanical load regulates bone remodeling through RANKL (Fu *et al*, 2002). The cellular source of RANKL for osteoclast formation under physiological and some pathological conditions like postmenopausal osteoporosis has not been fully revealed. Mice harboring a conditional knockout of RANKL using *Prx1^Cre^* demonstrate no osteoclast formation, which indicates that mesenchyme-derived cells are an essential source of RANKL in the development of long bones (Xiong *et al*., 2011). Among the mesenchyme-derived cells, matrix-embedded hypertrophic chondrocytes and osteocytes are the major source of RANKL for osteoclastogenesis, not osteoblasts or their progenitors. Hypertrophic chondrocytes control the resorption of calcified cartilage during bone development, while osteocytes regulate bone remodeling by sensing mechanical changes (Wang *et al*, 2020a; Xiong *et al*., 2011). And osteocytes are an essential source of RANKL for cancellous bone remodeling in adult mice but not younger mice (Nakashima *et al*., 2011; Xiong *et al*., 2011).

RANKL is produced in both soluble and membrane-bound forms. Since matrix-embedded cells have limited direct contact with osteoclast progenitors and in postmenopausal osteoporosis plasma soluble RANKL is significantly increased (Jabbar *et al*, 2011), it is proposed that the soluble RANKL is a major form to regulate osteoclastogenesis. Nevertheless, growing mice lacking soluble RANKL showed no deficits in bone mass and equal amounts of bone loss after estrogen deprivation compared to controls (Xiong *et al*., 2018). These studies indicate that membrane-bound RANKL is sufficient for bone growth and is responsible for bone loss induced by estrogen deficiency. Therefore, it is mysterious how matrix-embedded cell-derived RANKL regulates osteoclastogenesis and bone remodeling.

Although marrow adipose tissue accounts for almost 70% of the adult bone marrow volume in humans, the functions of BM adipose lineage cells are still very mysterious (Scheller & Rosen, 2014). Clinically, increased marrow adipose tissue is associated with increased fracture risk in diseases such as postmenopausal osteoporosis, anorexia nervosa and diabetes (Veldhuis-Vlug & Rosen, 2018). Mesenchyme-derived BM adipose lineage cells express RANKL, which is distinctive from other adipose tissues, including both white and brown adipose (Xiong *et al*., 2018). Udagawa et al. first explored that bone marrow-derived preadipose cell line MC3T3-G2/PA6 cells support osteoclast formation in the co-culture system (N. *et al*., 1989). This is the first hint of the regulatory effects of bone marrow derived preadipocytes on osteoclastogenesis. Several studies have explored the roles of bone marrow adipose lineage cells by using the *Adipoq^Cre^* (Ambrosi *et al*., 2017; Zhou *et al*., 2017). Zhou et al. reported that bone marrow adipose lineage cells promoted the regeneration of stem cells and hematopoiesis by secreting SCF (Zhou *et al*., 2017). The author demonstrated that four weeks after gavaging 6-week-old *Adipoq^Cre/ER^; R26^tdTomato^* mice with tamoxifen, tdTomato was expressed in 93 ± 5% of perilipin^+^ adipocytes. The tdTomato expressing in the bone marrow besides adipocytes was a subset of LepR^+^ stromal cells: 5.9 ± 3.1% of LepR^+^ cells in *Adipoq^Cre^*^/*ER*^*; R26^tdTomato^* bone marrow were tdTomato^+^. They concluded that *Adipoq^Cre^*^/*ER*^ recombines adipose lineage cells in the bone marrow. *Adipoq^Cre^*^/*ER*+^ cells rarely formed osteoblasts *in vivo*. In another study by Thomas et al., lineage tracing results showed that the *Adipoq^Cre^* targeted almost all bone marrow adipose lineage cells but not matrix-embedded cells, including hypertrophic chondrocytes or osteocytes (Ambrosi *et al*., 2017). A recent study by Leilei *et al*. (Zhong *et al*., 2020) identified a distinct population of non-lipid laden adipose-lineage cells targeted by the *Adipoq^Cre^* by employing the single cell transcriptome analysis. The lineage tracing of the *Adipoq^Cre^* demonstrated that at 1 month of age, Td labeled all perilipin^+^ adipocytes, CD45^−^ stromal cells with a reticular shape, and pericytes, but not osteoblasts, osteocytes, periosteal surface, growth plate or articular chondrocytes, which further confirmed that the *Adipoq^Cre^* rarely labels mesenchymal progenitors that generate osteoblasts, osteocytes and chondrocytes. *In vivo* transplantation under the kidney capsule showed that bone marrow Td^+^ cells from 1-month-old Col2/Td mice but not *Adipoq-Td* mice form bone-like structure. A recent study by Hisa *et al*. reported that the *Adipoq^Cre^* could efficiently target a subset group of mesenchymal stromal cells (Mukohira *et al*., 2019). The *Adipoq^Cre^* could efficiently target PDGFR β^+^VCAM-1^+^ and CXCL12-abundant reticular (CAR) stromal cells which are characterized as pericytes contributing to the vessel maintenance and hematopoietic niche. Lineage tracing showed that tdTomato expression was rarely detected in ALP^+^ osteoblasts of *Adipoq^Cre^; R26^tdTomato^* mice at 8 weeks old. BMSCs are a group of heterogenous stromal cells with various markers and functions. PDGFR β^+^ and CAR stromal cells represent bone marrow pericytes important for vascular functions (Andrae *et al*, 2008; Hellstrom *et al*, 1999). Interestingly, Leilei *et al*. reported that after adipocytes in 1-month-old *Adipoq^Cre^; Rosa-Tomato*/*DTR* (Adipoq/Td/DTR) mice was ablated via diphtheria toxin (DT) injections, BM vasculature became dilated and distorted, and the number of Emcn^+^CD31^+^ endothelial cells were decreased (Zhong *et al*., 2020). The results imply that the non-lipid laden adipocyte described by Leilei *et al*. is probably the same subgroup of mesenchymal stromal cells characterized by Hisa *et al*. Given the current evidence, we used the *Adipoq^Cre^* to explore the roles of BM adipocyte derived RANKL in osteoclastogenesis.

In our study, we found that loss of RANKL production by BM adipose lineage cells did not alter osteogenic or adipogenic differentiation of bone marrow mesenchymal stromal progenitors. Lineage tracing results showed the *Adipoq^Cre^* efficiently targeted BM adipose lineage cells and unlike the BMSCs, the tdTomato^+^ cells from *Adipoq^Cre^; R26^tdTomato^* mice mainly formed adipocytes and could not proliferate and undergo osteogenesis, chondrogenesis differentiation *in vitro* and *in vivo*. Based on the previous studies and our findings, we conclude that the *Adipoq^Cre^* could efficiently target the BM adipose lineage cells.

*Adipoq^Cre^; rankl^fl/fl^* mice showed increased numbers of trabeculae, elevated BMD, decreased osteoclast formation in the marrow, and decreased resorption of cancellous bone. The number of osteoblasts and bone formation in the distal femur were also reduced. The reduced osteogenesis was not due to the decreased BMSCs differentiation capacity, but secondary to decreased osteoclastogenesis and bone turnover rate. These results demonstrate that RANKL from BM adipose lineage cells contributes to trabecular bone remodeling in growing and adult mice. The difference in BMD between *Adipoq^Cre^; rankl^fl/fl^* and *rankl^fl/fl^* mice is smaller in adult mice than in growing mice. This is probably due to the increasing impact of RANKL from osteocytes on trabecular bone remodeling in adult mice.

TRAP^+^ cells in different areas could possess distinct functions. Resorption of calcified cartilage in the growth plates is essential for bone elongation (Lewinson & Silbermann, 1992). In growing mice, loss of BM adipocyte RANKL does not affect resorption of calcified cartilage. TRAP staining demonstrated the number of TRAP^+^ osteoclasts under the growth plate, identified as a non-bone-resorbing osteoclast subtype called vessel-associated osteoclast (VAO)(Romeo *et al*, 2019), was not altered after the deletion of RANKL. Periosteal TRAP^+^ cells could regulate cortical bone formation (Gao *et al*, 2019). Macrophage-lineage TRAP^+^ cells induced transcriptional expression of periostin and recruitment of periosteum-derived cells (PDCs) to the periosteal surface through secretion of PDGF-BB, where the recruited PDCs underwent osteoblast differentiation coupled with type H vessel formation, which is different from the trabecular bone remodeling. Loss of RANKL in BM adipose lineage cells did not affect cortical bone thickness or the number of periosteal TRAP^+^ cells.

The function of soluble RANKL on osteoclast formation and bone resorption has recently been identified. Mice lacking soluble RANKL show normal tooth eruption and bone development. Soluble RANKL contributes to the formation of osteoclasts in cancellous bone in adult mice, with membrane-bound RANKL proving sufficient for normal growth and skeletal development (Xiong *et al*., 2018). Our findings showed that the bone marrow soluble RANKL levels were not significantly different between *Adipoq^Cre^; rankl^fl/fl^* and *rankl^fl/fl^* mice, which indicated that BM adipose lineage cells are not an important source of sRANKL production. In some pathological conditions, including PMOP, periodontitis, and inflammatory joint disease, soluble RANKL levels are elevated (Chen *et al*, 2010; Geusens *et al*, 2006; Nile *et al*, 2013), which indicates that soluble RANKL is functionally involved in these conditions. However, mice lacking soluble RANKL showed a similar amount of bone loss after estrogen withdrawal (Xiong *et al*., 2018), which implies that soluble RANKL is not essential for osteoclasts formation and bone resorption after estrogen withdrawal. In our study, no bone loss was observed in *Adipoq^Cre^; rankl^fl/fl^* mice after estrogen withdrawal. Osteoclast formation and bone resorption were significantly inhibited following the deletion of RANKL in BM adipose lineage cells. Furthermore, the bone turnover rate was significantly lower in *Adipoq^Cre^; rankl^fl/fl^* mice compared to their littermate controls. These results suggest that RANKL from BM adipose lineage cells contributes to excess osteoclast formation and bone resorption in pathological bone loss diseases with increased BM adiposity.

ROS is widely clinically used to lower blood glucose levels in diabetic patients via the activation of PPARγ. However, the use of ROS is associated with an increased risk of fractures as well as an increase in BM adipocyte counts due to the activation of PPARγ, as essential mediator of adipocyte differentiation and function (Aubert *et al*., 2010; Kahn *et al*., 2008), Although the role of PPARγ in osteoclast differentiation is still a topic of considerable debate (Mbalaviele *et al*, 2000; Wan *et al*, 2007; Wei *et al*., 2010; Zou *et al*, 2016), our findings showed that administering ROS to *Adipoq^Cre^; rankl^fl/fl^* mice failed to increase osteoclast formation and bone resorption or induce bone loss, as observed in *rankl^fl/fl^* mice. These results indicate that ROS increases osteoclast formation and bone resorption at least in part by increasing the number of BM adipose lineage cells.

Osteocytes sense mechanical loading and secrete RANKL to control osteoclast formation and bone remodeling in adult mice and unloading-induced bone loss (Wang *et al*., 2020a; Xiong *et al*., 2011). Since the *Adipoq^Cre^* does not target osteocyte, we further testify whether adipocyte RANKL mediates skeletal mechanical loading. Tail-suspension did induce a significant loss of cancellous bone volume in both *Adipoq^Cre^; rankl^fl/fl^* and *rankl^fl/fl^* mice, indicating adipocyte RANKL did not mediate unloading-induced bone resorption.

Taken together, our study demonstrates that the RANKL from BM adipose lineage cells serves as an essential source of trabecular bone remodeling in both physiological and pathological conditions but does not mediate resorption of calcified cartilage in growth plate or regulate cortical bone remodeling.

## Materials and methods

### Mice and study design

Wild type (WT) C57BL/6J mice were purchased from vitalriver Company (Beijing, China). The *Adipoq^Cre^* mice were obtained as a generous gift from Prof. Ma Xinran from the East China Normal University. *Rankl^fl/fl^* and *rank^fl/fl^* mice were purchased from the Jackson Laboratory (JAX stocks No. 018978 and 027495). *Rosa26-lsl-tdTomato* mice were provided by the Shanghai Model Organisms (Shanghai, China). Cross-fertilization, breeding, and feeding of all mice were performed at Shanghai Model Organisms (SCXK [Shanghai] 2017-0010 and SYXK [Shanghai] 2017-0012). The mice were kept in SPF facilities with no more than 5 in a cage with food and water available ad libitum. The protocols are in compliance with regulations of the ethical committee of Shanghai Changhai Hospital. Mice were anesthetized at defined intervals (4, 8, and 16 weeks) by 5% chloral hydrate to collect femur, blood, centrum, bone marrow and tooth samples. Samples were divided into different groups randomly, and littermates were used as controls. Sample sizes were calculated on the assumption that a 30% difference in the parameters measured would be considered biologically significant with an estimate of sigma of 10–20% of the expected mean.

### *In vivo* pathological bone loss models

Female mice of 8 weeks age were anesthetized with 5% chloral hydrate and then ovariectomized via transection to establish the postmenopausal osteoporosis (PMOP) model. After 6 weeks, the mice were anesthetized by 5% chloral hydrate to collect femur and blood.

Female mice of 8 weeks age were treated with rosiglitazone (10 mg/kg/day, intragastric administration). After 6 weeks, the mice were anesthetized by 5% chloral hydrate to collect femur and blood.

For the unloading model, we performed the tail suspension. The tail of mice was tied to the rearing cage. After 4 weeks, the femur was collected and scanned by micro-CT.

### Cell culture

To verify the proliferation capacity of tdTomato^+^ cells of *Adipoq^Cre^; R26^tdTomato^* mice, tdTomato^+^ cells were sorted by flow cytometry. And bone marrow adipose lineage cells were collected following previously reported.(Fan *et al*., 2017) The tdTomato^+^ cells were cultured in DMEM medium (Hyclone, Logan, UT, USA) supplemented with 10 % fetal bovine serum (Hyclone), and the colony forming unit-fibroblast (CFU-F) was analyzed by the giemsa (Beyotime, Shanghai, China) staining after 7 days culture. Bone marrow mesenchymal stromal cells (BMSCs) were seeded onto 96-well plates and treated with dexamethasone (10^−8^ mol/L), β-glycerophosphate (10 mM), and ascorbic acid (50 mg/mL) for 3 days to induce osteogenesis. After 21 days, the alizarin red staining was performed to detect osteogenesis. BMSCs were induced by β-sodium glycerophosphate (10 mmol/L), dexamethasone (10^−8^ mol/L), and vitamin C (10 mmol/L) for adipogenesis.(Xin *et al*, 2018) Adipocyte formation was analyzed via oil red O staining.

### ELISA

Commercially available ELISA kits were used to determine concentrations of active CTX-1 (novus, catalog no. NBP2-69074), OCN (novus, catalog no. NBP2-68151), and RANKL (Abcam, catalog no. ab100749) in bone marrow according to the manufacturer’s instructions.

### Real-time PCR

The total RNA of cells was collected by RNA fast 200 kit (Catalog no. 220010, Feijie bio company, Shanghai, China). We amplified and detected *β-actin* (mouse: 5′-TCTGCTGGAAGGTGGACAGT-3′ [forward], 5′-CCTCTATGCCAACACAGTGC-3′ [reverse]), *Rankl* (mouse: 5′-TGTACTTTCGAGCGCAGATG-3′ [forward], 5′-AGGCTTGTTTCATCCTCCTG-3′ [reverse]), *PPARγ* (mouse: 5′-GGAAGACCACTCGCATTCCTT-3′ [forward], 5′-GTAATCAGCAACCATTGGGTCA-3′ [reverse]), *Lpl* (mouse: 5′-ATGGATGGACGGTAACGGGAA-3′ [forward], 5′-CCCGATACAACCAGTCTACTACA-3′ [reverse]), *Alp* (mouse: 5′-CCAACTCTTTTGTGCCAGAGA-3′ [forward], 5′-GGCTACATTGGTGTTGAGCTTTT-3′ [reverse]) and *Ocn* (mouse: 5′-GGACCATCTTTCTGCTCACTCTGC-3′ [forward],

5′-TGTTCACTACCTTATTGCCCTCCTG-3′ [reverse]) using a Real-Time PCR system (analytic Jena), as described previously.(Chen *et al*, 2018b)

### Western blotting

Protein expression was determined by Western blotting using standard methods. Blots were probed with anti-LPL (ab91600, 1:500), anti-PPARγ (ab59256, 1:800), anti-OCN (ab93876, 1:500), anti-ALP (ab229126, 1:1000), and anti-RANKL (ab45039, 1:300) antibodies (Abcam, Cambridge, Britain), as described previously.(Luo *et al*, 2016; Ma *et al*, 2017)

### Flow cytometry

Cells were collected from femurs of *rankl^fl/fl^* mice, *Adipoq^Cre^; rankl^fl/fl^* mice, *Adipoq^Cre^; R26^tdTomato^* mice, after which bone marrow cells were collected and labeled with monoclonal anti-mouse CD3 (Sigma, catalog no. SAB4700050, 1: 1000), CD19 (Sigma, catalog no. SAB4700108, 1: 1000) antibodies, CD45 (eBioscience, catalog no. 11-0451, 1: 500), CD31 (BD Bioscience, catalog no. 558738, 1: 500), Sca-1 (eBioscience, catalog no. 56-5981, 1: 500), PDGFα (Biolegend, catalog no. 135907, 1: 500) and Reep2 (Invitrogen, catalog no. MA5-25878, 1: 500 and Abcam, catalog no. ab6563, 1: 1000). Cells were analyzed on an LSRII flow cytometer system (BD Biosciences) using FlowJo software (Tree Star), as described previously.(Chen *et al*., 2018a) tdTomato^+^ cells were sorted and collected from the femur of *Adipoq^Cre^; R26^tdTomato^* mice.

### *In vivo* heterotopic bone formation assay

The sorted tdTomato^+^ cells from *Adipoq^Cre^; R26^tdTomato^* mice and BMSCs were collected and 0.5 × 10^6^ cells mixed with 40 mg hydroxyapatite in 100 μl α-MEM and incubated overnight. Then the cells were implanted under the kidney capsules of BALB/c nude mouse for 4 weeks. After retrieval, it was paraformaldehyde-fixed and paraffin-embedded. The slice was stained with OCN, Aggrecan, Perilipin and positive cells were counted.

### Micro-CT and X ray

Bone was fixed with 4% paraformaldehyde for 24 h and analyzed by Skyscan 1172 high-resolution micro-CT (Skyscan, Antwerp, Belgium). Scan conditions were set at 8 µm per pixel resolution, 80 kV voltage, and 124 μA current. Using these images, we constructed three-dimensional models and analyzed images by CTAn and CTVol, including bone mineral density (BMD), bone volume/tissue volume (BV/TV), trabecular number (Tb.N), trabecular separation (Tb.Sp), and cortical thickness. And the tooth eruption was evaluated by the X ray.

### Histochemistry, immunohistochemistry, and immunofluorescent staining

Femurs were fixed in 4% paraformaldehyde for 24 h and decalcified in 10% EDTA (room temperature) for 2 weeks. After dehydration in an alcohol gradient (50%, 75%, 85%, 95%, or 100%), femurs were embedded in paraffin, cut into 4 µm thick sections using a Thermo Scientific Microm HM 325 Rotary Microtome, and processed for hematoxylin-eosin (H&E) staining and TRAP staining according the manufacturer’s instructions (Collins *et al*, 2017). For immunohistochemistry staining, sections were incubated with primary antibodies against RANKL (Abcam; catalog no. ab216484, 1:100) with a horseradish peroxidase-conjugated anti-rabbit secondary antibody used to detect immunoactivity. For immunofluorescence staining, frozen sections were incubated with primary antibodies overnight to osteocalcin (Biocompare; catalog no. bs-4917R; 1:100), adiponectin (Abcam; catalog no. ab181281; 1:200), aggrecan (proteintech; catalog no. 13880-1-AP; 1:200) and perilipin (Invitrogen; catalog no. PA1-1051; 1:100). Then the sections were incubated with corresponding fluorescence-conjugated secondary antibodies (1:200) and DAPI (1:500) for 1 h at room temperature. Then the samples were scanned on either a Nano Zoomer or 3D Histech system, with exported images analyzed using Case Viewer and NDP view software.

### Calcein staining

To detect mineral deposition, we injected mice with alizarin red S (Sigma; 40 mg/kg body weight) 10 days and calcein (Sigma; 20 mg/kg body weight) 3 days before euthanasia. We performed undecalcified bone slicing and calculated the mineral apposition rate (MAR) of trabecular bone and cortical bone.

### Statistics

All of the analysis were performed by an independent investigator blinded to the genotypes of the animals under analysis. Data are presented as means ± SDs. Independent-samples *t* tests and one-way ANOVAs were used to assess statistical significance. All analyses were performed using SPSS 21.0 (IBM, Chicago, IL, USA). Statistical significance was set at *P* < 0.05.

## Data availability

This study includes no data deposited in external repositories.

## Acknowledgments

We thank Prof. Ma Xinran from the East China Normal University for giving us the *Adipoq^Cre^* mice as a generous gift. All the transgenic mice in the study were bred and housed in Shanghai Model Organisms Center Inc. We thank the *Textcheck* for language polishing. The English in this document has been checked by at least two professional editors, both native speakers of English. For a certificate, please see: http://www.textcheck.com/certificate/zWIVZE. This work was supported by the National Key (2018YFC2001500); National Natural Science Foundation (NNSF) Key Research Program in Aging (91749204); National Natural Science Foundation of China (81871099, 81771491); Municipal Human Resources Development Program for Outstanding Leaders in Medical Disciplines in Shanghai (2017BR011); Shanghai Young Physician Supporting Program (2018.15).

## Author Contributions

XC, YH, XZ and JCS: study design. XZ, WC, HWC, XQL, YJW, LPW, JC, YH, LHC, WC, YHL, YL, JWG, SCW and WZW: data collection. XZ, CF, QRZ, YJW, BTH and XQL: data analysis. XC and JCS: data interpretation. XC and XZ: drafting manuscript. XC and JCS: revising manuscript content. JCS: approving final version of manuscript.

## Conflict of interests

The authors declare no conflict of interests.

---

**Figure EV1.**
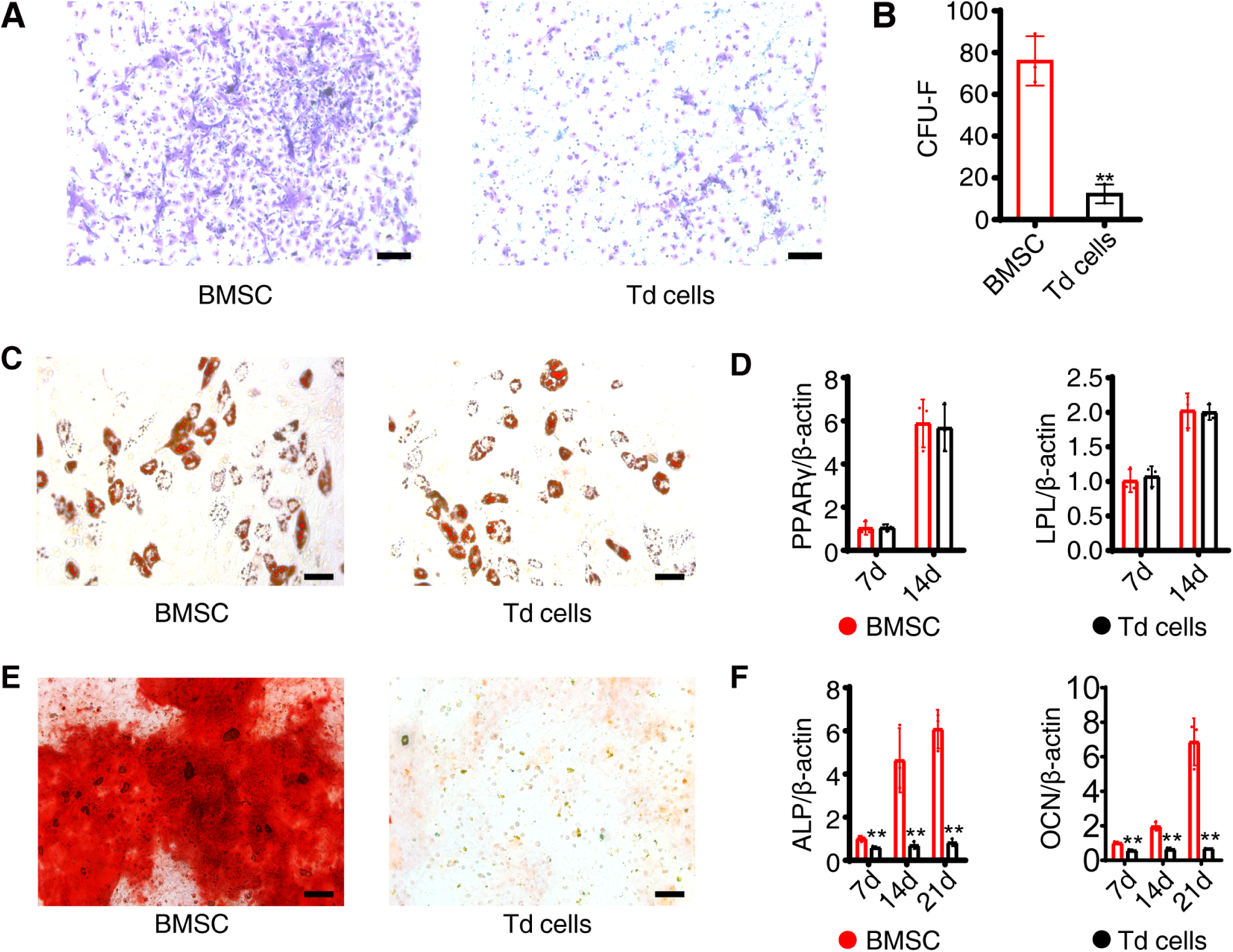
CFU-F, adipogenic and osteogenic differentiation analysis *in vitro*. (A-B) Giemsa staining and statistical analysis of CFU-F number of BMSCs from WT mice and tdTomato^+^ cells from *Adipoq^Cre^; R26^tdTomato^* mice. Scale bar=50 µm. Data were compared using an unpaired t-test (** indicates *P* < 0.01), error bars are standard deviations. n=5 independent microscopic vision fields in (B). (C-D) Adipogenic potential of BMSCs and tdTomato^+^ cells were assessed by oil red O staining and *PPARγ* and *Lpl* were detected by qPCR. Scale bar=50 µm. Data were compared using an unpaired t - test, error bars are standard deviations. n=5 independent microscopic vision fields in (D). (E-F) Osteogenic potential of BMSCs and tdTomato^+^ cells were assessed by the alizarin red staining and *Alp* and *Ocn* were detected by qPCR. Scale bar=50 µm. Data were compared using an unpaired t-test (** indicates *P* < 0.01), error bars are standard deviations. n=5 independent microscopic vision fields in (F).

**Figure EV2.**
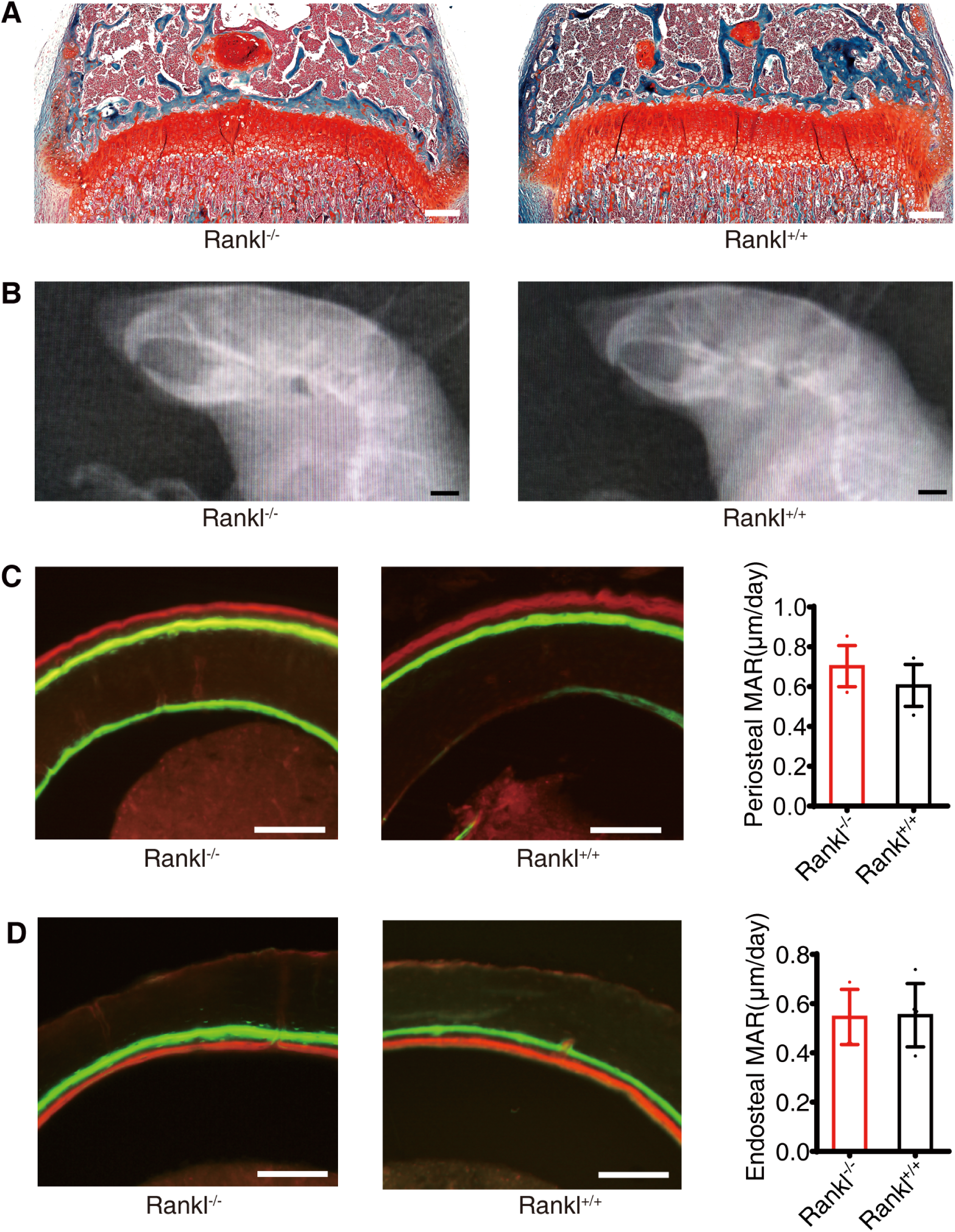
Adipogenic, osteogenic and chondrogenic differentiation analysis *in vivo*. (A) Immunofluorescence staining of OCN and statistical analyses of BMSCs and tdTomato^+^ cells (Td cells), scale bar=50 µm. Images are representative of five independent biological replicates, n=6 independent microscopic vision fields in the right panel. Data were compared using an unpaired t-test (** indicates *P* < 0.01), error bars are standard deviations. (B) Immunofluorescence staining of Aggrecan and statistical analyses of positive cells, scale bar=50 µm. Images are representative of five independent biological replicates, n=6 independent microscopic vision fields in the right panel. Data were compared using an unpaired t-test (** indicates *P* < 0.01), error bars are standard deviations. (C) Immunofluorescence staining of Perilipin and statistical analyses of positive cells, scale bar=50 µm. Images are representative of five independent biological replicates, n=6 independent microscopic vision fields in the right panel. Data were compared using an unpaired t-test (** indicates *P* < 0.01), error bars are standard deviations.

**Figure EV3.**
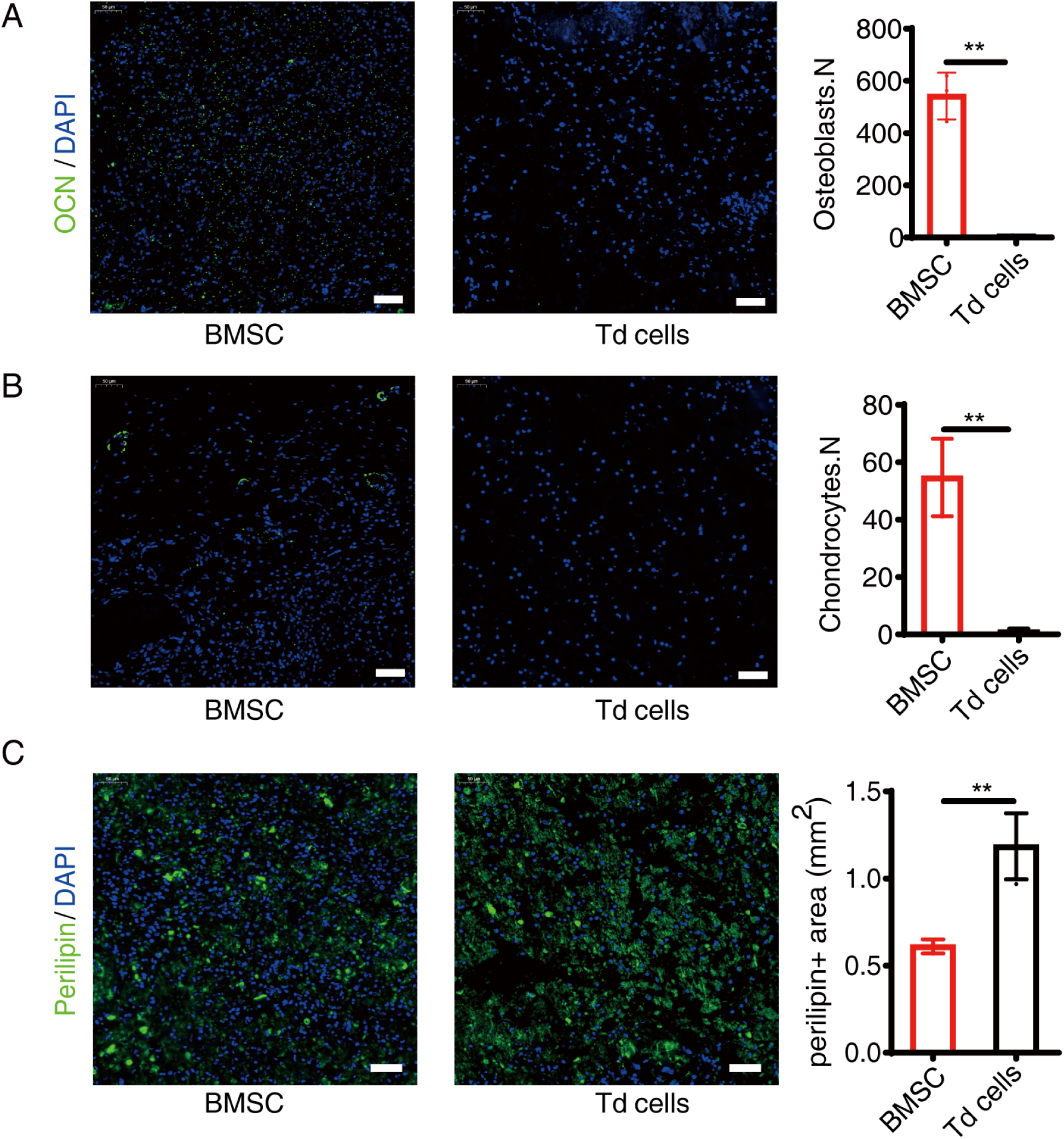
RANK deletion in BM adipose lineage cells does not affect bone mass. (A) Safranin O/Fast green staining of *Adipoq^Cre^; rankl^fl/fl^* (*Rankl^-/-^*) and *rankl^fl/fl^* (*Rankl^+/+^*) mice. Scale bar=250 µm. (B) X ray images of the head of 8-week-old mice. Scale bar= 2mm. (C) Calcein staining and statistical analysis from periosteal side of cortical bone (n=6 mice in *Rankl*^-/-^ group and 5 mice in *Rankl*^+/+^ group). Scale bar=50 μm. Data were compared using an unpaired t-test, error bars are standard deviations. (D) Calcein staining and statistical analysis from endosteal side of cortical bone (n=6 mice in *Rankl*^-/-^ group and 5 mice in *Rankl*^+/+^ group). Scale bar=50 μm. Data were compared using an unpaired t-test, error bars are standard deviations.

**Figure EV4.**
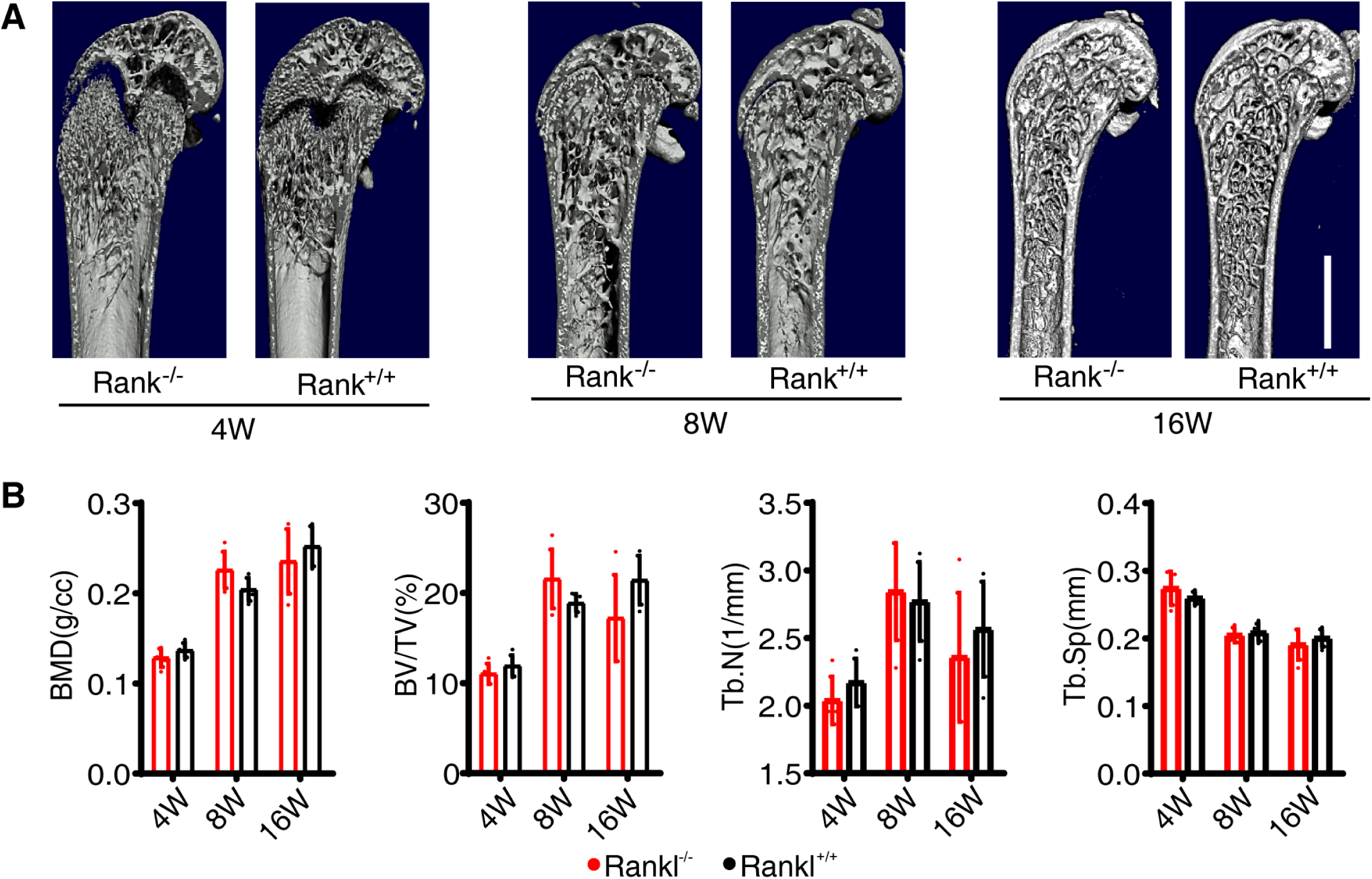
RANK deletion in BM adipose lineage cells does not affect bone mass. (A-B) Micro-CT and statistical analyses of BMD, BV/TV, Tb.N and Tb.Sp in trabecular bone from *Adipoq^Cre^; rank^fl/fl^* (*Rank^-/-^*) and *rank^fl/fl^* (*Rank^+/+^*) mice at 4 weeks, 8 weeks and 16 weeks. Scale bar=1 mm. Images are representative of 5 independent biological replicates, n=5 independent microscopic vision fields in (B). Data were compared using an unpaired t-test, error bars are standard deviations.

**Figure EV5.**
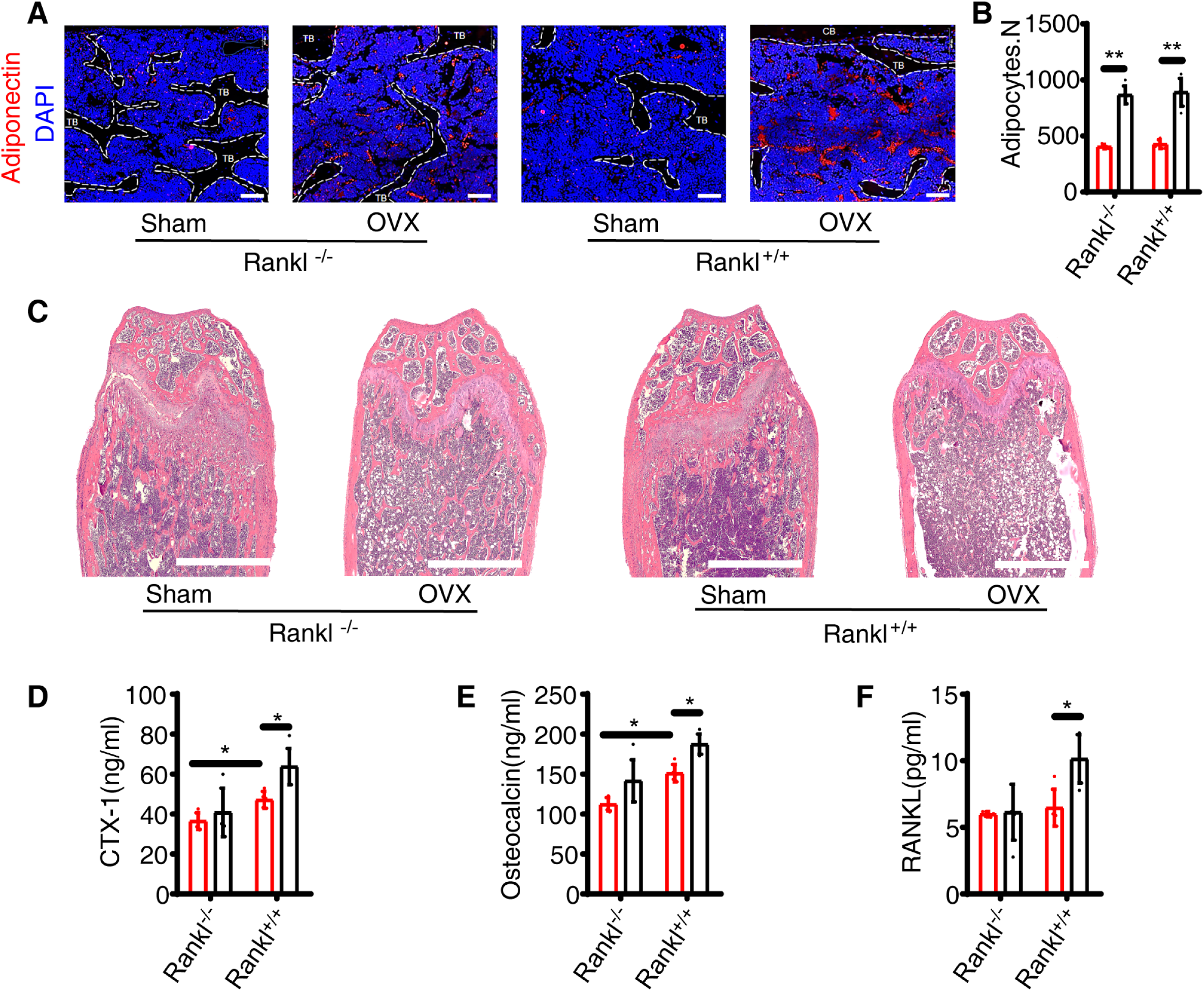
RANKL knockout from BM adipose lineage cells contributes to OVX-induced bone loss. (A-B) Immunofluorescence staining of adiponectin and statistical analyses of the number of adipose lineage cells. Scale bar=100 µm. Images are representative of 5 independent biological replicates, n=5 independent microscopic vision fields in (B). Data were compared using an unpaired t-test (** indicates *P* < 0.01), error bars are standard deviations. CB: cortical bone, TB: trabecular bone. (C) H&E staining of the distal femur from *Adipoq^Cre^; rankl^fl/fl^* (*Rankl^-/-^*) and *rankl^fl/fl^* (*Rankl^+/+^*) mice treated with Sham or OVX. Scale bar=1 mm. (D-F) ELISA analyses of serum CTX-1, OCN and bone marrow RANKL levels in Sham and OVX mice (n=5 independent biological replicates). Data were compared using an unpaired t-test (* indicates *P* < 0.05), error bars are standard deviations.

## Notes

### Competing Interest Statement

The authors have declared no competing interest.

## References

Ambrosi TH, Scialdone A, Graja A, Gohlke S, Jank AM, Bocian C, Woelk L, Fan H, Logan DW, Schurmann A et al (2017) Adipocyte Accumulation in the Bone Marrow during Obesity and Aging Impairs Stem Cell-Based Hematopoietic and Bone Regeneration. Cell stem cell 20: 771–784.e776

Andrae J, Gallini R, Betsholtz C (2008) Role of platelet-derived growth factors in physiology and medicine. Genes Dev 22: 1276–1312

Aubert RE, Herrera V, Chen W, Haffner SM, Pendergrass M (2010) Rosiglitazone and pioglitazone increase fracture risk in women and men with type 2 diabetes. Diabetes Obes Metab 12: 716–721

Boyle WJ, Simonet WS, Lacey DL (2003) Osteoclast differentiation and activation. Nature 423: 337–342

Chen CH, Chen HA, Liao HT, Liu CH, Tsai CY, Chou CT (2010) Soluble receptor activator of nuclear factor-kappaB ligand (RANKL) and osteoprotegerin in ankylosing spondylitis: OPG is associated with poor physical mobility and reflects systemic inflammation. Clinical rheumatology 29: 1155–1161

Chen X, Zhi X, Wang J, Su J (2018a) RANKL signaling in bone marrow mesenchymal stem cells negatively regulates osteoblastic bone formation. Bone research 6: 34

Chen X, Zhi X, Yin Z, Li X, Qin L, Qiu Z, Su J (2018b) 18beta-Glycyrrhetinic Acid Inhibits Osteoclastogenesis In Vivo and In Vitro by Blocking RANKL-Mediated RANK-TRAF6 Interactions and NF-kappaB and MAPK Signaling Pathways. Frontiers in pharmacology 9: 647

Collins FL, Williams JO, Bloom AC, Singh RK, Jordan L, Stone MD, McCabe LR, Wang ECY, Williams AS (2017) CCL3 and MMP-9 are induced by TL1A during death receptor 3 (TNFRSF25)-dependent osteoclast function and systemic bone loss. Bone 97: 94–104

Duque G, Li W, Adams M, Xu S, Phipps R (2011) Effects of risedronate on bone marrow adipocytes in postmenopausal women. Osteoporosis international : a journal established as result of cooperation between the European Foundation for Osteoporosis and the National Osteoporosis Foundation of the USA 22: 1547–1553

Fan Y, Hanai JI, Le PT, Bi R, Maridas D, DeMambro V, Figueroa CA, Kir S, Zhou X, Mannstadt M et al (2017) Parathyroid Hormone Directs Bone Marrow Mesenchymal Cell Fate. Cell Metab 25: 661–672

Fu Q, Jilka RL, Manolagas SC, O’Brien CA (2002) Parathyroid hormone stimulates receptor activator of NFkappa B ligand and inhibits osteoprotegerin expression via protein kinase A activation of cAMP-response element-binding protein. J Biol Chem 277: 48868–48875

Gao B, Deng R, Chai Y, Chen H, Hu B, Wang X, Zhu S, Cao Y, Ni S, Wan M et al (2019) Macrophage-lineage TRAP+ cells recruit periosteum-derived cells for periosteal osteogenesis and regeneration. J Clin Invest 129: 2578–2594

Geusens PP, Landewe RB, Garnero P, Chen D, Dunstan CR, Lems WF, Stinissen P, van der Heijde DM, van der Linden S, Boers M (2006) The ratio of circulating osteoprotegerin to RANKL in early rheumatoid arthritis predicts later joint destruction. Arthritis and rheumatism 54: 1772–1777

Hellstrom M, Kalen M, Lindahl P, Abramsson A, Betsholtz C (1999) Role of PDGF-B and PDGFR-beta in recruitment of vascular smooth muscle cells and pericytes during embryonic blood vessel formation in the mouse. Development (Cambridge, England) 126: 3047–3055

Jabbar S, Drury J, Fordham JN, Datta HK, Francis RM, Tuck SP (2011) Osteoprotegerin, RANKL and bone turnover in postmenopausal osteoporosis. 64: 354–357

Kahn SE, Zinman B, Lachin JM, Haffner SM, Herman WH, Holman RR, Kravitz BG, Yu D, Heise MA, Aftring RP et al (2008) Rosiglitazone-associated fractures in type 2 diabetes: an Analysis from A Diabetes Outcome Progression Trial (ADOPT). Diabetes care 31: 845–851

Kronenberg HM (2003) Developmental regulation of the growth plate. Nature 423: 332–336

Lacey DL, Timms E, Tan HL, Kelley MJ, Dunstan CR, Burgess T, Elliott R, Colombero A, Elliott G, Scully S et al (1998) Osteoprotegerin ligand is a cytokine that regulates osteoclast differentiation and activation. Cell 93: 165–176

Lewinson D, Silbermann M (1992) Chondroclasts and endothelial cells collaborate in the process of cartilage resorption. The Anatomical record 233: 504–514

Liu WG, Toyosawa S, Furuichi T, Kanatani N, Yoshida C, Liu Y, Himeno M, Narai S, Yamaguchi A, Komori T (2001) Overexpression of Cbfa1 in osteoblasts inhibits osteoblast maturation and causes osteopenia with multiple fractures. Journal of Cell Biology 155: 157–166

Luo J, Yang Z, Ma Y, Yue Z, Lin H, Qu G, Huang J, Dai W, Li C, Zheng C et al (2016) LGR4 is a receptor for RANKL and negatively regulates osteoclast differentiation and bone resorption. Nat Med 22: 539–546

Ma YM, Fu ST, Lu L, Wang XH (2017) Role of androgen receptor on cyclic mechanical stretch-regulated proliferation of C2C12 myoblasts and its upstream signals: IGF-1-mediated PI3K/Akt and MAPKs pathways. Molecular and Cellular Endocrinology 450: 83–93

Manolagas SC, Jilka RL (1995) Bone marrow, cytokines, and bone remodeling. Emerging insights into the pathophysiology of osteoporosis. N Engl J Med 332: 305–311

Mbalaviele G, Abu-Amer Y, Meng A, Jaiswal R, Beck S, Pittenger MF, Thiede MA, Marshak DR (2000) Activation of peroxisome proliferator-activated receptor-gamma pathway inhibits osteoclast differentiation. J Biol Chem 275: 14388–14393

Mukohira H, Hara T, Abe S, Tani-Ichi S, Sehara-Fujisawa A, Nagasawa T, Tobe K, Ikuta K (2019) Mesenchymal stromal cells in bone marrow express adiponectin and are efficiently targeted by an adiponectin promoter-driven Cre transgene. International immunology 31: 729–742

N. U, N. T, T. A, T. S, A. Y, H. K, T.J. M, T. S (1989) The bone marrow-derived stromal cell lines MC3T3-G2/PA6 and ST2 support osteoclast-like cell differentiation in cocultures with mouse spleen cells. Endocrinology 125: 1805–1813

Nakashima T, Hayashi M, Fukunaga T, Kurata K, Oh-Hora M, Feng JQ, Bonewald LF, Kodama T, Wutz A, Wagner EF et al (2011) Evidence for osteocyte regulation of bone homeostasis through RANKL expression. Nature Medicine 17: 1231–1234

Nile CJ, Sherrabeh S, Ramage G, Lappin DF (2013) Comparison of circulating tumour necrosis factor superfamily cytokines in periodontitis patients undergoing supportive therapy: a case-controlled cross-sectional study comparing smokers and non-smokers in health and disease. J Clin Periodontol 40: 875–882

Raggatt LJ, Partridge NC (2010) Cellular and molecular mechanisms of bone remodeling. J Biol Chem 285: 25103–25108

Romeo SG, Alawi KM, Rodrigues J, Singh A, Kusumbe AP, Ramasamy SK (2019) Endothelial proteolytic activity and interaction with non-resorbing osteoclasts mediate bone elongation. Nature cell biology 21: 430–441

Scheller EL, Rosen CJ (2014) What’s the matter with MAT? Marrow adipose tissue, metabolism, and skeletal health. Ann N Y Acad Sci 1311: 14–30

Sobacchi C, Frattini A, Guerrini MM, Abinun M, Pangrazio A, Susani L, Bredius R, Mancini G, Cant A, Bishop N et al (2007) Osteoclast-poor human osteopetrosis due to mutations in the gene encoding RANKL. Nat Genet 39: 960–962

Veldhuis-Vlug AG, Rosen CJ (2018) Clinical implications of bone marrow adiposity. J Intern Med 283: 121–139

Wan Y, Chong LW, Evans RM (2007) PPAR-gamma regulates osteoclastogenesis in mice. Nat Med 13: 1496–1503

Wang L, You X, Lotinun S, Zhang L, Wu N, Zou W (2020a) Mechanical sensing protein PIEZO1 regulates bone homeostasis via osteoblast-osteoclast crosstalk. Nat Commun 11: 282

Wang Y, Wang Y, Chen Y, Qin Q (2020b) Unique epidemiological and clinical features of the emerging 2019 novel coronavirus pneumonia (COVID-19) implicate special control measures. Journal of medical virology 92: 568–576

Wei W, Wang X, Yang M, Smith LC, Dechow PC, Sonoda J, Evans RM, Wan Y (2010) PGC1beta mediates PPARgamma activation of osteoclastogenesis and rosiglitazone-induced bone loss. Cell Metab 11: 503–516

Xin Z, Jin C, Chao L, Zheng Z, Liehu C, Panpan P, Weizong W, Xiao Z, Qingjie Z, Honggang H et al (2018) A Matrine Derivative M54 Suppresses Osteoclastogenesis and Prevents Ovariectomy-Induced Bone Loss by Targeting Ribosomal Protein S5. Frontiers in Pharmacology 9

Xiong J, Cawley K, Piemontese M, Fujiwara Y, Zhao H, Goellner JJ, O’Brien CA (2018) Soluble RANKL contributes to osteoclast formation in adult mice but not ovariectomy-induced bone loss. Nat Commun 9: 2909

Xiong J, Onal M, Jilka RL, Weinstein RS, Manolagas SC, O’Brien CA (2011) Matrix-embedded cells control osteoclast formation. Nat Med 17: 1235–1241

Yu B, Huo L, Liu Y, Deng P, Szymanski J, Li J, Luo X, Hong C, Lin J, Wang CY (2018) PGC-1alpha Controls Skeletal Stem Cell Fate and Bone-Fat Balance in Osteoporosis and Skeletal Aging by Inducing TAZ. Cell stem cell 23: 615–623

Zhong L, Yao L, Tower RJ, Wei Y, Miao Z, Park J, Shrestha R, Wang L, Yu W, Holdreith N et al (2020) Single cell transcriptomics identifies a unique adipose lineage cell population that regulates bone marrow environment. eLife 9

Zhou BO, Yu H, Yue R, Zhao Z, Rios JJ, Naveiras O, Morrison SJ (2017) Bone marrow adipocytes promote the regeneration of stem cells and haematopoiesis by secreting SCF. Nat Cell Biol 19: 891–903

Zhou BO, Yue R, Murphy MM, Peyer JG, Morrison SJ (2014) Leptin-receptor-expressing mesenchymal stromal cells represent the main source of bone formed by adult bone marrow. Cell Stem Cell 15: 154–168

Zou W, Rohatgi N, Chen TH, Schilling J, Abu-Amer Y, Teitelbaum SL (2016) PPAR-gamma regulates pharmacological but not physiological or pathological osteoclast formation. Nat Med 22: 1203–1205

